# Combined inhibition of S100A4 and TIGIT suppresses late-stage breast cancer metastasis to the lung by activating T and NK cells

**DOI:** 10.1101/2024.11.24.624743

**Authors:** Jia-Shiun Leu, Hui Deng, Han Nhat Tran, Xiong Wei, Chihiro Hashiomoto, Fransisca Leonard, Thomas Wong, Nourhan Abdelfattah, José M Benítez Salazar, Reece Kang, Jose A Maldonado, Carlo D. Cristobal, Hyun Kyoung Lee, Zhiqiang An, Ningyan Zhang, Kyuson Yun

**Affiliations:** Department of Neurology, Houston Methodist Research Institute, Houston, TX, USA; Texas Therapeutics Institute, Brown Foundation Institute of Molecular Medicine, The University of Texas Health Science Center at Houston, Houston, TX 77030, USA; Department of Pediatrics, Baylor College of Medicine, Houston, TX 77030, USA; Duncan Neurological Research Institute, Texas Children’s Hospital, Houston, TX 77030, USA; Department of Neurology, Weill Cornell Medical College, New York, NY 10021, USA

## Abstract

Cancer metastasis is responsible for approximately 90% of cancer-related deaths, but very few treatment options exist currently. While the role of S100A4 in promoting metastasis has been known for decades, this knowledge has not been translated in the clinic. Here, we report that a novel monoclonal antibody against the human S100A4 protein (S1004-11) effectively suppresses breast cancer metastasis in two different mouse models. Importantly, a novel combination of ant-TIGIT and S100A4-11 can suppress lung metastases even in late-stage disease after the lung premetastatic niche (PMN) has already been established similar to a stage when many breast cancer patients are diagnosed. Mechanistically, S1004-11 mAb treatment block the formation of PMN by suppressing neutrophil infiltration and activating natural killer (NK) and T cells in the lung. Single-cell RNA-sequencing and cell:cell communication analyses indicate that TIGIT signaling suppresses NK cells, which is reversed in S100A4-11 treated PMN. In summary, this study provides compelling evidence for a novel mechanism of S100A4 function in PMN formation and the feasibility of using S100A4-11 monoclonal antibody to suppress metastases at different stages of breast cancer progression.

## Text main Introduction

Cancer is a major cause of death worldwide, accounting for 10 million deaths in 2020 (*1*). Due to earlier detection and effective treatment, overall survival for some cancers, such as breast and colon cancer, has increased significantly in the last decade. However, only limited progress has been made to treat metastatic diseases which accounts for ∼ 90% of cancer-related deaths (*2, 3*). During the metastatic process, primary tumor cells must leave the primary tumor site, intravasate into blood vessels, survive and migrate in the circulation, extravasate into secondary tissue, and colonize and proliferate in target organs. For successful colonization, tumor cells need to overcome short nutrient supply and escape host immune surveillance. To facilitate this process, cancer cells prime a distinct “soil” microenvironment before they “seed” the metastatic site (*4*).

Recent studies suggest that immune cells play critical roles in promoting metastasis, especially myeloid cells from the bone marrow. Both monocytic myeloid derived suppressor cells (M-MDSCs) and granulocytic/polymorphic nuclear myeloid derived suppressor cells (PMN-MDSCs) and neutrophils have been implicated in promoting cancer metastasis (*5, 6*). For example, neutrophils recruited to the lung have been shown to generate pre-metastatic niches (PMN) PMNs for breast cancer cells (*7–10*): depletion of neutrophils (and/or PMN-MDSCs) with anti-Ly6G antibody treatment decreased lung metastasis (*8, 10*). In addition, neutralizing antibody treatments for G-CSF (*11*) or GM-CSF (*9, 10*), chemokines for neutrophils, dramatically reduced the number of lung metastatic nodules in treated mice. G-CSF elimination in a spontaneous breast metastatic model also inhibited cancer metastasis (*8*), and a known mechanism of neutrophils in the PMNs is have suppressing cytotoxic T and NK cells from eliminating cancer cells (*12, 13*). These and other studies firmly established that PMN formation by myeloid cell infiltration, particularly granulocytes, is an important step that may be targeted for therapeutic benefit.

S100A4, a member of the S100 protein family and also known as metastasin and fibroblast specific protein1 (*FSP1*), has been studied as a mediator of breast cancer metastasis for decades. It promotes epithelial-mesenchymal transition, angiogenesis, cancer metastasis and stemness in multiple cancer types, including breast (*14*), brain (*15, 16*), and colorectal cancer (*17, 18*). Earlier studies have shown that genetic deletion of *S100a4* in the MMTV-PyMT breast cancer mouse model reduced lung metastasis (*19*). Likewise, a blocking antibody against the mouse S100A4 protein suppressed lung metastasis but not primary tumor growth (*20–22*). However, mechanisms through which S100A4 promotes metastasis are still not fully understood and whether S100A4 plays a role in late stages of disease progression. Notably, deletion of *S100a4* only in the tumor microenvironment (TME) is sufficient to significantly extend survival in glioma mouse models (*15*), suggest that S100A4 inhibition in the TME may be a viable approach to suppress cancer metastasis and tumor progression.

T-cell immunoreceptor with immunoglobulin and ITIM domain (TIGIT) is an immune-checkpoint molecule expressed on exhausted CD8^+^ T cells, NK cells, and regulatory T (Treg) cells. TIGIT ligands CD155 (PVR) and CD112 (PVRL2) are expressed on tumor and antigen-present cells within TME (*23*) and suppresses T and NK cell function. TIGIT is often co-expressed with programmed death-1 (PD-1) and cytotoxic T lymphocyte-associated protein 4 (CTLA4) on exhausted T cells, as well as on NK cells. These co-expression patterns may explain why ant-PD-1 therapy alone is insufficient in many cancer types (*24*). Due to these differential expression patterns of immune checkpoint molecules, it is postulated that combining TIGIT with other immunotherapies may more effectively restore anti-tumor immune responses in the TME (*25, 26*). Here, we demonstrate that *S100a4* in the host is necessary for the E0771 breast cancer mouse allograft growth. Additionally, a blocking monoclonal antibody against the human S100A4 protein, S100A4-11, suppresses lung metastases in the MMTV-PyMT and 4T1 breast tumor mouse models. Mechanistically, S1004-11 antibody treatment blocks neutrophil infiltration to generate PMNs and reactivation of cytotoxic NK and T cells in the lung at both early and late stages metastases. As such, a novel combination therapy consisting of anti-TIGIT and S100A4-11 significantly suppresses lung metastases even during late-stage breast tumor progression.

## RESULTS

### Deletion of *S100a4* in stromal cells is sufficient to block breast tumor growth

Previously, we reported that S100A4 confers immune suppressive phenotypes in tumor infiltrating T and myeloid cells in human brain cancer, and that deletion of *S100a4* in the host mouse was sufficient to significant extend survival of glioma-bearing mice (*15*). To test whether *S100a4* plays a similar tumor-promoting role in other cancer types, we transplanted E0771 murine breast cancer cells into mammary fat pads of *S100a4*^-/-^, *S100a4^+^*^/-^ or B6 wildtype control mice. Surprisingly, no breast tumors formed in *S100a4*^-/-^ host mice, even at day 21 after mammary fat-pad injection (**Figure 1a**). Furthermore, E0771 tumor growth was significantly reduced in *S100a4*^+/-^ mice compared to wildtype control host mice. These results indicate that S100A4 plays a critical role in tumor promoting stromal cells for multiple cancer types.

**Fig 1.**
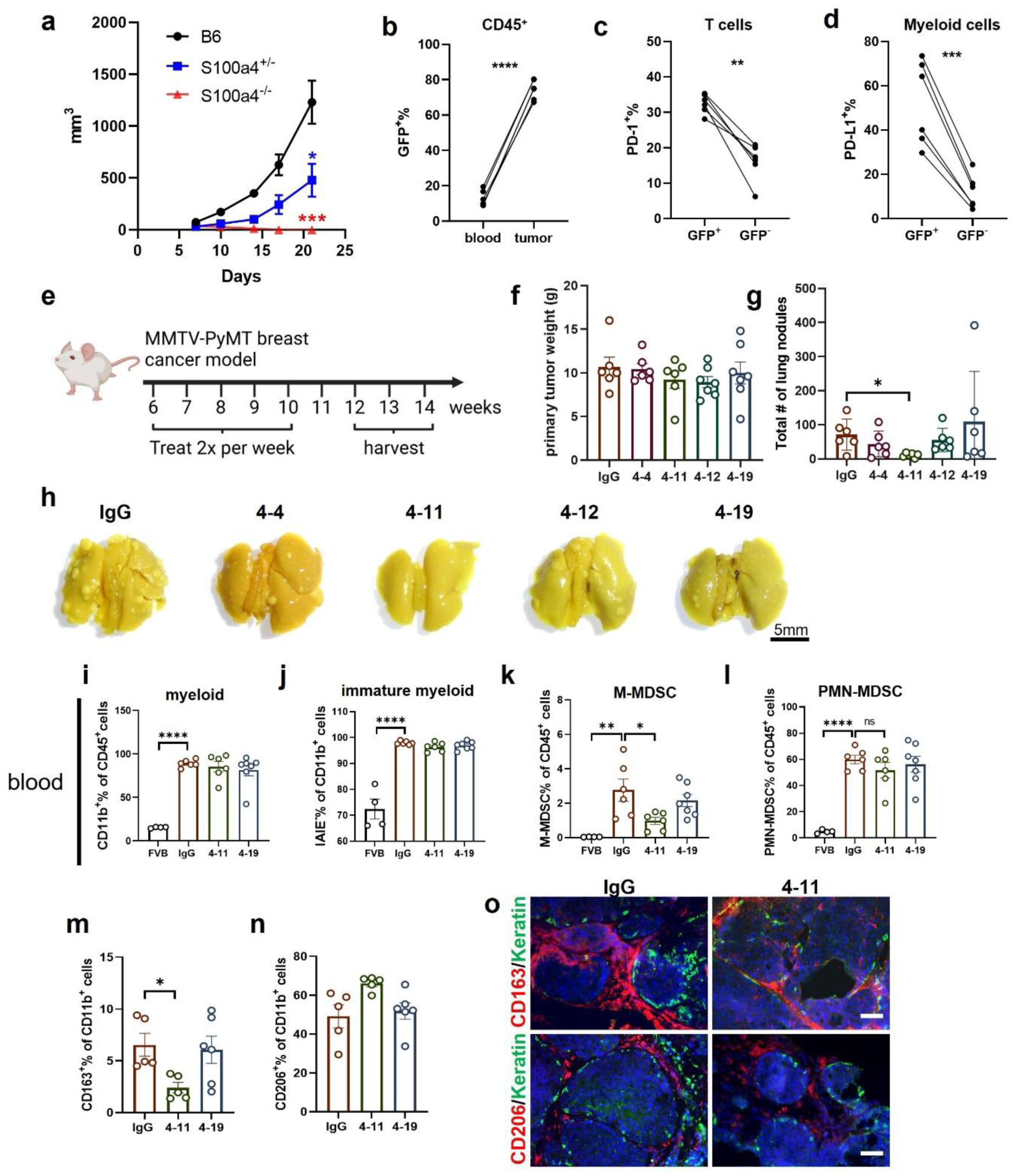
S100A4-11 antibody suppresses spontaneous lung metastases in MMTV-PyMT mouse model. (a) The mammary fat pads of B6, *S100a4^+/-^*, *S100a4^-/-^* mice were injected with 5×10^5^ E0771 cells (R2). Tumor volume was measured twice a week for 3 weeks (n=5-6). Multiple *t*-tests was used. \**p*<0.05; *** *p*<0.001. (b) GFP^+^ expression level in the immune cells are measured in the E0771 tumor and blood. Paired *t-*test analysis was used. \*\*\*\**p*<0.0001. (c) PD-1^+^ frequency of GFP^+^ or GFP^-^ cells from the T cells are measured in the E0771 tumor of *S100a4^+/-^* mice (n=6). Paired *t-* test analysis was used. ** *p*<0.01. (d) PD-L1^+^ frequency of GFP^+^ or GFP^-^ cells from myeloid cells are measured in the E0771 tumor of *S100a4^+/-^* mice (n=6). Paired *t-*test analysis was used. *** *p*<0.001. (e) 6-week-old MMTV-PyMT transgenic female mice were treated with IgG control or one of four different monoclonal antibodies against S100A4, twice per week for 4 weeks. The mice were harvested when the total tumor burden reached a humane endpoint (12-14 weeks of age). (f, g) The total number of primary tumor weight and lung nodules were measured at harvest (n=6-7). Kruskal-Wallis test was applied. (h) Representative images of lungs from different treatment groups fixed in Bouin’s solution. Scale bar denotes 5 mm. (i-l) Flow cytometry analysis of myeloid cells (i), immature myeloid cells (j), and M-MDSCs (k), and PMM-MDSC (l) in the blood from FVB control (n=4) and MMTV-PyMT (n=6-7 per group). (m, n) Flow cytometry analysis of CD163^+^ and CD206^+^ macrophages in breast tumors of S100A4-11 and S100A4-19 antibody-treated mice, compared with IgG control (n=5-6). (o) Immunofluorescence analysis of breast tumor-associated CD163^+^ and CD206^+^ macrophage from MMTV-PyMT mice. The scale bar denotes 50µm. (i-n) Ordinary one-way ANOVA analysis was used when comparing groups. Error bar indicates mean ± SEM. * *p*<0.05; ** *p*<0.01; **** *p*<0.0001.

To understand which cell types express S100A4 in the stroma, we leveraged the *S100a4*^+/-^ mice which has the green fluorescence protein (GFP) cDNA knocked-into the *S100a4* locus (*27*). We previously validated that the endogenous S100A4 expression is faithfully recapitulated by GFP expression in this mouse (*16*). Therefore, we used flow cytometry to determine which immune cell types express S100A4 in the TME and blood of *S100a4*^+/-^ mice bearing E0771 breast tumor in the mammary fat pad. GFP^+^/S100a4^+^ cells constituted <20% (range from 9-19.5%) of all immune cells in the blood but >60% (range from 67.3-80.4%) of all immune cells in the TME (**Figure 1b**). Overall, GFP^+^/S100a4^+^ expression is increased in the tumor-associated T and myeloid cells (**Supplementary Figure 1a**,1c). Among CD11b^+^PDL1^+^ myeloid cells, ∼45% (range 34.7-54%) are GFP^+^/S100a4^+^ in the blood but nearly all are GFP^+^/*S100a4*^+^ in the TME (range 96.3-98.1%) (**Supplementary Figure 1b**, n=6 paired samples). Similarly, among CD3^+^ PD1^+^ T cells, 5-30% in the blood are GFP^+^/*S100a4*^+^ while 49-77.3% of them are GFP^+^/*S100a4*^+^ in the TME (**Supplementary Figure 1d**, n=6 paired samples). Conversely, when GFP^+^ and GFP^-^ T cells in the TME were further analyzed, ∼30% (range from 28.1-35.3%) of GFP^+^ T cells were PD1^+^ and while <20% (range from 6.22-20.8%) of GFP^-^ T cells were PD1^+^, indicating that S100A4^+^ are significantly enriched in the immune suppressive T cells (**Fig. 1c,** n=6). In the myeloid cells, GFP^+^/*S100a4*^+^ cells are also significantly higher in immune suppressive myeloid cells (CD11b^+^PDL1^+^) than GFP^-^ cells (**Fig. 1d,** n=6). These results strongly support targeting S100A4 in the TME to reverse immune suppression in breast cancer.

### A novel S100A4 blocking antibody generation and characterization

To translate our findings, we developed and characterized a S100A4 blocking antibody to inhibit S100A4 function in the TME. Rabbits were immunized with the recombinant human S100A4 protein, and a total of 114 B cell clones were screened (**Supplementary Figure 2a**). To screen the monoclonal antibodies against S100A4, we adapted the chick DRG neurite outgrowth assay based on an earlier study that showed that the soluble S100A4 protein induced neurite growth of astrocytes and DRG from chick embryos (*28*). We tested 114 clones in triplicates. Neurite outgrowth was significantly inhibited after pre-incubating the S100A4 protein with S100A4 -1, -5, or -11 monoclonal antibody (**Supplementary** Fig. 2b-2c). The variable regions of these and a few other select clones were cloned and sequenced. Then, full-length IgG1 antibodies were expressed and purified from HEK293 cultures. The binding affinity of each clone was determined by biolayer interferometry (BLI), and the kinetic binding (KD) of S100A4-1, -9, -11, and -19 antibodies were found in the nanomolar range (**Supplementary Figure 2d**).

To determine its pharmacokinetic properties, S100A4-11 clone was injected into B6 females (20mg/kg) and various organs and blood were analyzed with ELISA for S100A4-11 levels. S100A4-11 antibody is very stable and persists in the serum and various organs. At 4 hours post injection, the maximum concentration in the blood was 100µg/ml and other organs were 10-20µg/g tissue, except in the brain (0.2-0.5 µg/g) (**Supplementary Figure 3a**-3f). The serum level at 7 days (168 hours) post injection remained at ∼80% of initial concentration. In other organs, including the lung, S100A4-11 remained at ∼10µg/g, >50% of maximum concentration at 7 days, indicating its stability.

Next, we confirmed the sequence similarity for the S100A4 protein and evaluated cross-species binding by top two clones, S100A4-11 and S100A4-19. S100A4 is a highly conserved protein (*29*), and its amino acids sequence is approximately 96% identical between human and mouse (**Supplementary Figure 2e**). We measured the affinity of S100A4-11 and S100A4-19 to both human and mouse S100A4 proteins by ELISA. S100A4-11 and -19 showed strong binding to both human and mouse S100A4 proteins (**Supplementary Figure 2f**-2g), enabling us to perform *in vivo* characterization of these antibodies in mouse models.

### S100A4-11 treatment suppresses lung metastases in MMTV-PyMT mice

MMTV-PyMT is a spontaneous breast cancer metastasis mouse model with a well characterized disease progression timeline and mimics the disease progression in human (*30*); therefore, we used this breast cancer model to evaluate the efficacy of our top S100A4 blocking monoclonal antibodies *in* vivo. MMTV-PyMT females (6-week-old) were intravenously injected with S100A4-4, -11, -12, -19 monoclonal antibodies or isotype control IgG antibodies (10mg/kg, twice/week for 4 weeks). The mice were euthanized once they reached humane end points due to mammary tumor burden (**Figure 1e**). Antibody treatments were well-tolerated, and no changes in body weight were detected during treatment (**Supplementary Figure 3g**). At end points, we measured both the combined weight of all primary mammary tumors and the total number of lung nodules from each mouse (**Figure 1f-1g**). While the primary tumor burden was equivalent in all groups (**Figure 1f**), S1004-11 mAb significantly reduced the number of lung nodules (**Figure 1g-1h**). Immunohistochemical analysis of lung nodules with an antibody against polyomavirus middle T antigen confirmed that lung nodules arose from MMTV-PyMT mammary tumors (**Supplementary Figure 3h**). Together, these results indicate that S1004-11 antibody is effective in suppressing spontaneous lung metastasis in the MMTV-PyMT model.

### S100A4-11 treatment reduces M-MDSCs and CD163+ macrophages

To elucidate potential mechanisms of action, flow cytometry was used to analyze the blood, mammary tumor, and metastatic nodule samples from mice treated with S100A4-11 or IgG control antibody. In the blood, MMTV-PyMT tumor bearing mice have significantly higher frequency of myeloid cells (CD11b^+^) than non-tumor bearing FVB mice (**Figure 1i**). Tumor-bearing mice also have higher frequency of immature myeloid cells (CD11b^+^/IAIE/MHCII^-^), also known as myeloid derived suppressive cells (MDSCs) (F**igure 1j**) than FVB control mice, consistent with previous reports (*31, 32*). S100A4-11 treatment significantly decreased the frequency of monocytic MDSCs (M-MDSC: CD11b^+^IAIE^-^Ly6C^hi^Ly6G^lo^, *p*=0.0483) (**Figure 1k**), but not polymorphonuclear MDSCs (PMN-MDSC: CD11b^+^IAIE^-^Ly6C^int^Ly6G^+^) (**Figure 1l**) in the blood. In contrast, no significant difference was detected in the number of infiltrating total immune cells (CD45^+^), myeloid cells (CD11b^+^), M-MDSC, and PMN-MDSCs in primary breast tumors (**Supplementary Figure 4a**-d).

To test if the changes in the immune cells in the blood of S100A4-11 treated mice alter the number of tumor infiltrating macrophages (TAMs), the frequency of immune-suppressive macrophages (CD163^+^, CD206^+^) were measured in the primary breast tumor (*31, 33, 34*). Flow cytometry analysis showed a significantly decreased frequency of CD163^+^ macrophages, but not CD206^+^ macrophages (**Figure 1m-1n**), in breast tumors of the S1004-11 antibody-treated mice, compared with IgG treated. Fixed tissue analysis with double-immunofluorescence confirmed reduced CD163^+^, but not CD206^+^, macrophages in S1004-11 antibody-treated MMTV-PyMT breast tumors (**Figure 1o**). Together, these results suggest that S100A4-11 antibody treatment reduces lung metastases, the number of circulating M-MDSCs, and immune suppressive CD163^+^ macrophages in the primary tumors of MMTV-PyMT mice.

### S100A4-11 treatment suppresses lung metastases in the 4T1 breast tumor model

To evaluate the generalizability of our findings, the efficacy of S1004-11 mAb was tested in another independent model of breast cancer metastasis in a different genetic background (BABL/c). The 4T1 allograft model mimics the triple-negative breast cancer and metastasizes to the lung and brain (*35*). BALB/c females were orthotopically injected with 4T1 cells in their mammary fat pad and treated intraperitoneally with either S1004-11 monoclonal or isotype control antibody (10 mg/kg, once a week), starting day 3 post-injection (**Figure 2a**). All mice were euthanized on 29 days post injection for analyses. The number of lung nodules was significantly reduced in the S1004-11 antibody-treated mice, compared with IgG antibody-treated mice (**Figure 2b, 2d**). Consistent with MMTV-PyMT model, S100A4-11 treatment did not affect primary/mammary tumor growth (**Figure 2c, Supplementary Figure 5a**), and the treatment was well tolerated in this genetic background as well (**Supplementary Figure 3i**).

**Fig 2.**
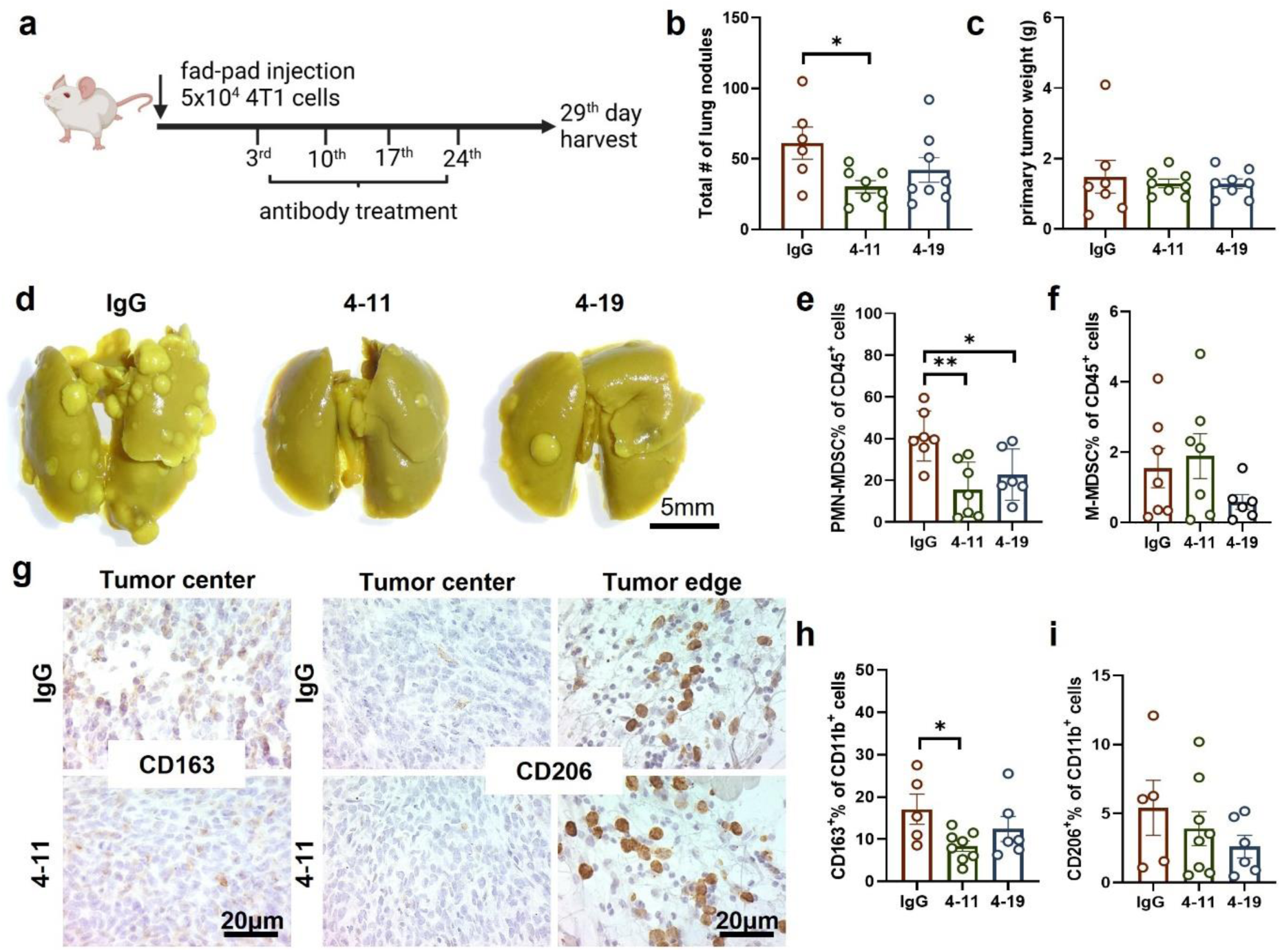
**S100A4-11 antibody treatment suppresses lung metastasis in the 4T1 breast cancer model**(a) The mammary fat pads of BALB/c female mice were injected with 5×10^4^ 4T1 cells for 3 days to establish breast tumors. The mice were then treated with IgG or S100A4 antibodies once a week for 4 weeks. (b, c) On day 29, the total number of lung nodules (b) and primary tumor weight (c) were recorded (n=6-8). The Kruskal-Wallis test was applied. (d) Representative lungs with metastatic nodules from IgG, 4-11, and 4-19 antibody-treated mice. The scale bar denotes 5 mm. (e, f) Flow analysis of lung samples show a significant decrease in PMN-MDSCs in both 4-11 and 4-19 antibody-treated mice. However, there is no significant change in M-MDSCs. (n=6-7) (g) Immunohistochemical analysis with CD163 and CD206 antibodies showing tumor center and edges in the 4T1 breast tumor (n=2). Scale bar denotes 20 µm. (h, i) Flow cytometry analysis of breast tumor tissues for CD163^+^ and CD206^+^ macrophages (n=5-8). (e, f, h, i) Ordinary one-way ANOVA analysis was used. Error bars denote mean ± SEM. *\*p<*0.05; *\*\*p<*0.01.

Like MMTV-PyMT mice, the frequencies of myeloid cells and MSDCs were higher in the blood of tumor bearing mice compared to wildtype control (**Supplementary Figure 5b**-5f). However, MDSC frequencies were not significantly reduced in the blood of S100A4-11 antibody-treated mice compared to control mice in this model. On the other hand, the frequency of PMN-MDSCs was significantly reduced in lung nodules of S100A4-11 antibody-treated mice, compared with isotype control antibody-treated mice (**Figure 2e)**. In contrast, no obvious difference was observed in the frequency of all immune infiltrates, myeloid cells, nor M-MDSCs in lung nodules (**Figure 2f, Supplementary Figure 5g, h).**

Next, we tested whether S100A4-11 treatment affected TAM frequency in the primary tumors. Immunohistochemical analysis showed that CD163^+^ and CD206^+^ macrophages preferentially localized to different tumor regions. Specifically, CD163^+^ macrophages were enriched in the tumor center, whereas CD206^+^ macrophages were enriched in tumor stromal region (**Figure 2g**), similar to other reports (*36*). Flow cytometry analyses reveal that the frequency of CD163^+^ TAMs was significantly reduced while the frequency of CD206+ TAMs was not affected in the S100A4-11 antibody-treated mice (**Fig. 2h, i)**, aligned with the observations in MMTV-PyMT model. There were no significant differences in the frequencies of immature myeloid cells and MDSCs in the breast tumor (**Supplementary Figure 5i**-l). Collectively, these findings indicate that in two independent breast tumor models, S100A4-11 antibody treatment reduces lung metastases and CD163^+^ TAM infiltration in breast tumors, supporting its broad efficacy in blocking breast cancer metastasis across different genetic backgrounds and tumor subtypes.

### S100A4-11 treatment blocks PMN formation in the lung by suppressing neutrophil infiltration

To investigate the mechanism of S100A4-11 efficacy in blocking lung metastases, multi-modal analyses were performed on lungs isolated from 4T1 tumor-bearing mice at an early timepoint (day 14 post-injection) when no detectable macro-metastasis (>2 mm) was present (*37*). First, to validate that circulating tumor cells are present at this point, fluorescently labeled 4T1 cells were injected into the mammary fat pads of the Balb/cJ females, and blood samples were collected at day 7 and day 14 post 4T1 inoculation in the mammary fat pad. We used flow cytometry to detect circulating 4T1 cells on day 7 post-injection and confirmed that their frequency is increased by day 14 (**Figure 3a**). In another cohort, 4T1 tumor-bearing mice were treated with either S100A4-11 or IgG isotype control antibody on days 3 and 10 post-injection. The mice were euthanized on day 14 post-injection, and the lungs were isolated and prepared for histology, RNA, and protein analyses (**Figure 3b**). Flow cytometry analysis revealed a marked increase in the frequency of Ly6G^+^ PMN-MDSCs/neutrophils in the lungs of the 4T1 mammary tumor-bearing mice at this timepoint, compared with non-tumor bearing BALB/cJ control mice (**Figure 3c**). Additionally, qPCR analyses of lung tissues showed that *S100a8* and *S100a9*, but not *S100a4* (p=0.1023), RNA expression is significantly elevated in the lungs of tumor bearing mice at day 14 post injection (**Figure 3d-3f**). This was consistent with a previous report that showed increased S100A8/9^+^ neutrophil infiltration in lungs of breast tumor bearing mice (*38*). Significantly reduced Ly6G^+^ neutrophil frequency (**Figure 3g)** as well as *S100a8/9/4* expression (**Figure 3h–3j**) were detected in the S1004-11 antibody-treated lungs compared to IgG control treated lungs, indicating that blocking S100A4 function significantly dampened the recruitment of neutrophils to the lungs.

**Fig 3.**
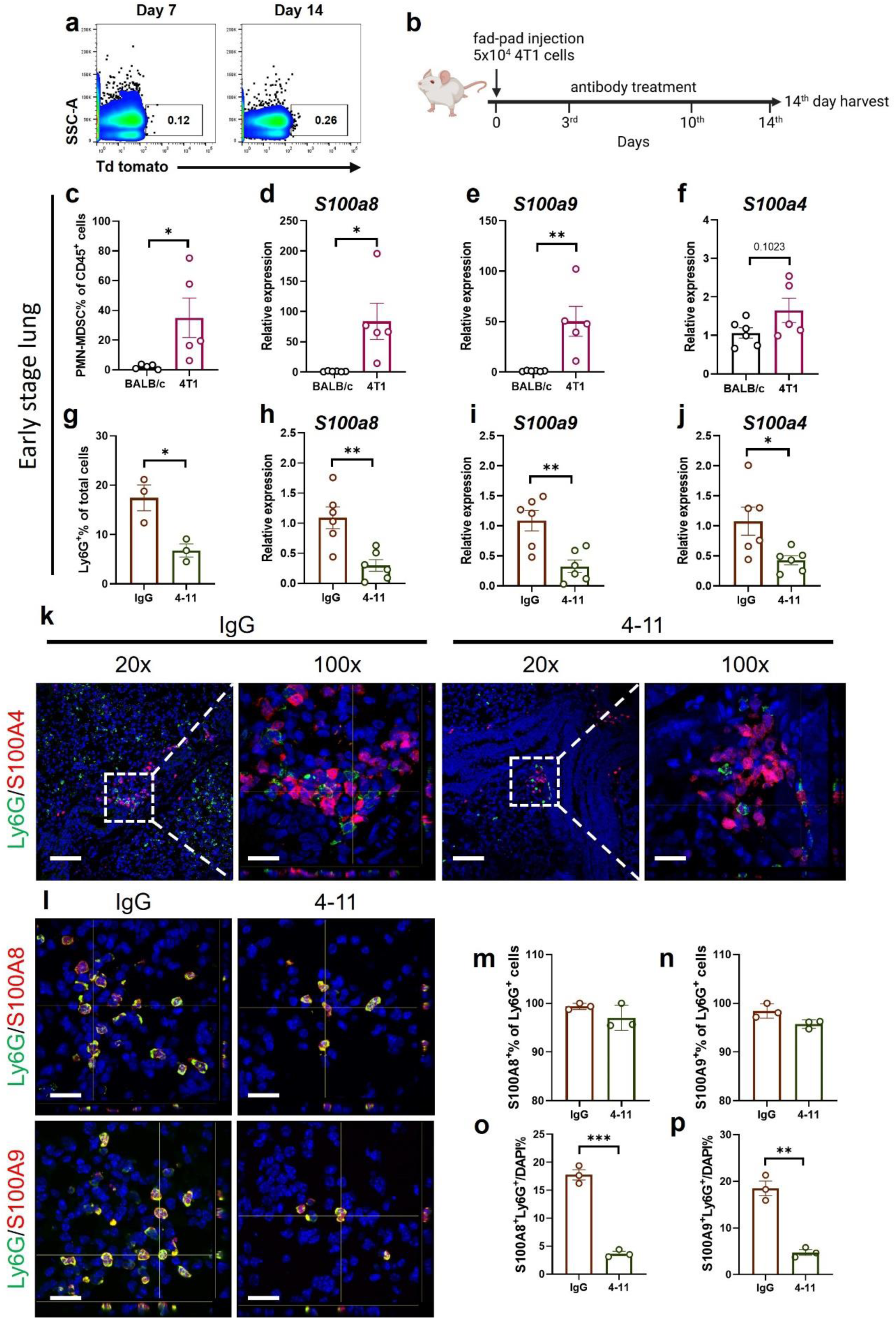
**S100A4-11 treatment blocks pre-metastatic niche formation in the lung**(a) Flow cytometry analysis shows the circulating Td tomato+ 4T1 cells are present in the blood on day 7 and day 14 post inoculation. (b) The mammary fat pad (R2) of BALB/c female mice were injected with 5 x 10^4^ 4T1 cells for 3 days to establish breast tumors. The mice were treated with IgG or anti-S100A4 antibodies on day 3 and day 10 before lungs were harvested on day 14. (c) Flow cytometry comparison of PMN-MDSCs in the lungs of 4T1 tumor- vs. non-tumor-bearing BALB/c mice (n=5 each). (d-f) *S100a8, S100a9, and S100a4* mRNA expression levels in tumor bearing vs. control mouse, measured by RT-PCR (n=5). (g-j) Comparisons of lung samples from IgG vs S100A4-11-antibody treated mice at day 14 post-injection. (f) Quantification of Ly6G+ cells in the lung measured by immunofluorescence image analysis (n=3). (g) RNA expression levels, measured by qPCR, of *S100a8* (g), *S100a9* (h), *and S100a4* (i) were significantly reduced in the 4-11 antibody-treated lung, compared to IgG treated group (n=6). (j) Representative confocal microscopy images showing Ly6G^+^ and S100A4^+^ cells. Scale bars denote 100 µm (20x) and 20 µm (100x). (k) Ly6G^+^ granulocytes co-express S100A8 and S100A9. Scale bar denotes 20 µm. (l, m) Quantification of immunofluorescence staining showing that >95% of S100A8 (l) or S100A9 (m) cells are Ly6G^+^ granulocytes in both IgG and 4-11 antibody-treated groups. (o, p) S100A8^+^Ly6G^+^ (o) and S100A9^+^Ly6G^+^ (p) cells are significantly decreased in the 4-11 treated lungs, compared with IgG control (n=3). Three representative fields/mouse lung were counted and averaged. Datapoints represent average per mouse per group. Unpaired *t-*test was used. Error bar means ± SEM. *\*p<*0.05; *\*\*p<*0.01.

To better define the changes in the premetastatic niche (PMN) at the cellular level, we performed double-immunofluorescence analyses with antibodies against S100A4, S100A8, S100A9, and Ly6G. S100A4^+^ and Ly6G^+^ cells are mutually exclusive (**Figure 3k**); however, >95% Ly6G+ cells co-express with S100A8 and S100A9 proteins in both S1004-11 and IgG antibody-treated lungs (**Figure 3l-3n**). The lung of S1004-11 antibody-treated mice contained a significantly reduced number of S100A8/9^+^/Ly6G^+^ neutrophils, compared to the isotype control antibody-treated mice (**Figure 3l, 3o-3p**). Altogether, these results indicate that soluble S100A4 protein promotes neutrophil (S100A8/9^+^/Ly6G^+^) infiltration into the lung to prime the PMN. Blocking S100A4 function with the S100A4-11 antibody significantly compromises the recruitment of these cells, thereby suppressing PMN formation.

### S1004-11 treatment reduces cytokines/chemokine production and MMP9 expression in the lung

To elucidate molecular consequences of S100A4-11 antibody treatment and blocked PMN formation, a targeted proteomics analysis was performed on the lungs (day 14 post-injection) of 4T1 tumor-bearing mice. C6 mouse cytokine arrays (RayBiotech) were used to measure the levels of 97 secreted protein in the lung lysates from IgG control or S100A4-11 treated 4T1 tumor bearing mice (**Supplementary Figure 6a**). Compared to isotype IgG treated mice, 10 proteins, including IL-20, MIP-1 gamma (CCL9), MIP-3 alpha (CCL20), Lungkine/CXCL15, osteopontin (OPN), pro-MMP9, SCF, TACI, TPO, and VEGFR1, were significantly reduced in the lungs of S1004-11 antibody-treated mice (**Figure 4a, Supplementary Figure 6b**). Notably, CCL9, CCL20, and CXCL15 are known chemokines that promote neutrophil infiltration (*39–41*) while IL-20 suppresses key inflammatory functions of neutrophils such as phagocytosis, granule exocytosis, and migration (*42*). Additionally, MMP9 has a critical role in regulating tumor vasculature remodeling and lung metastasis (*43, 44*). Double-immunofluorescence analysis confirmed that the majority of MMP9^+^ cells are also Ly6G^+^ (**Figure 4b-4c**) and they are significantly reduced in the lung from S1004-11 antibody-treated mice (**Figure 4b, 4d**). Further analyses by qPCR (**Figure 4e**) and western blot (**Figure 4f)** confirmed that MMP9 expression is reduced in the lung from S100A4-11 treated mice. Together, these results show that S100A4-11 antibody treatment reduced the production of chemokines and cytokines as well as the expression of TME remodeling factor (MMP9) critical for PMN formation and metastatic breast cancer cell colonization.

**Fig 4.**
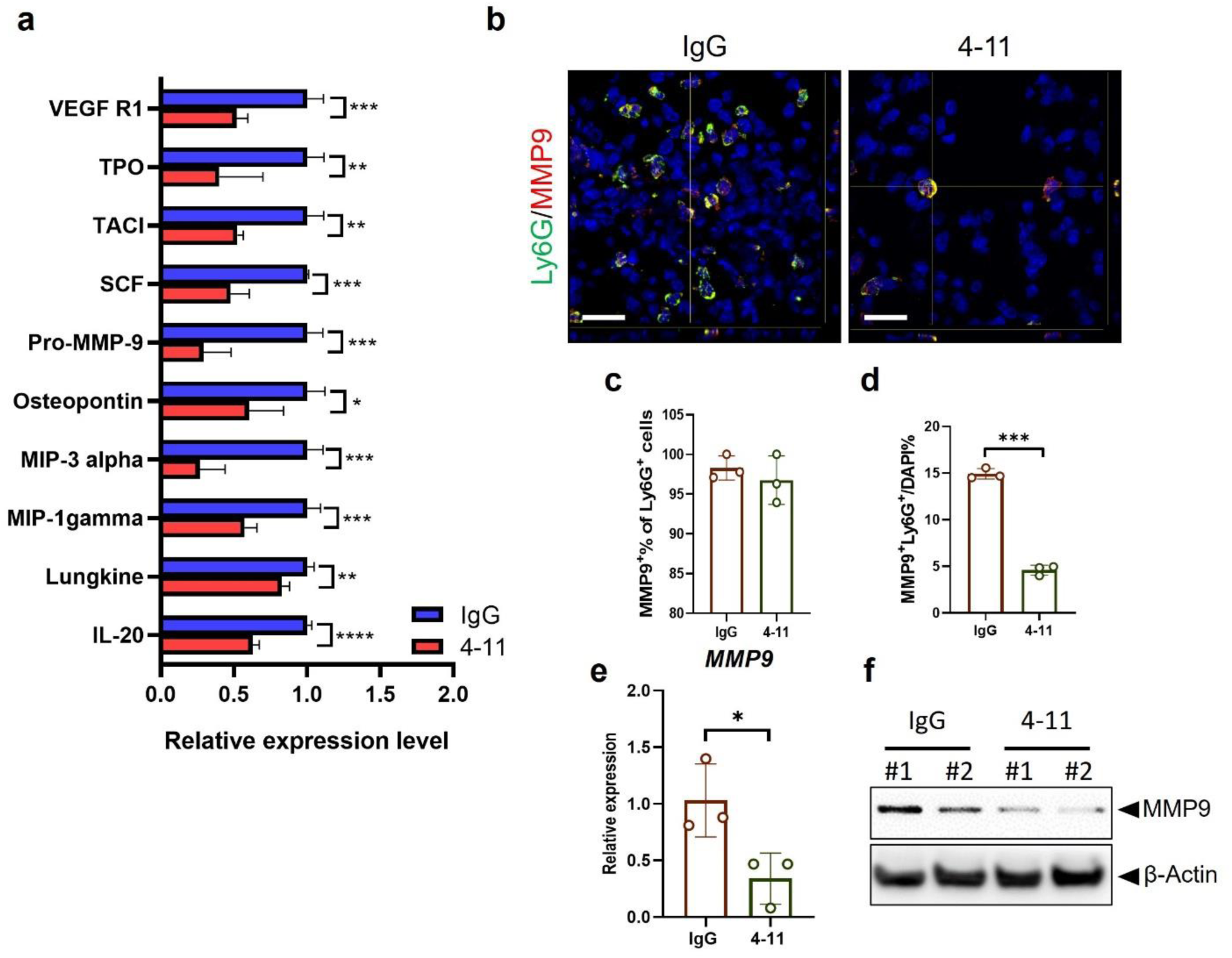
**S1004-11 treatment alters chemokine and cytokine expression in the pre-metastatic lung** (a) Lung lysates (day 14) from 4T1 tumor-bearing mice were analyzed by a cytokine array (n=2). 10 proteins were significantly reduced in the S100A4-11 antibody-treated lungs. Unpaired *t-*test was used. Error bar means ± SEM. *\*p<*0.05; *\*\*p<*0.01; *\*\*\*p<*0.001. (b) Representative images from double-immunofluorescence analysis of MMP9 and Ly6G in the lung. Scale bar denotes 20 µm. (c) Quantitation of MMP9^+^ cells as a percentage in Ly6G^+^ cells and (d) fractions of double-positive cells from all cells in the lungs of IgG and 4-11 treated mice. Each group was counted from three mice and three representative fields from each paraffin slide per mouse. Data shows averaged numbers per mouse per group. Analysis of MMP9 RNA (e) and protein (f) levels, measured by qPCR and western blotting respectively, in IgG and 4-11 treated lung at day 14. Unpaired *t-*test was used. Error bar is mean ± SEM. *\*p<*0.05; *\*\*\*p<*0.001

### S100A4-11 treatment increases lymphocyte infiltration and activation in the PMN

To gain more granular mechanistic understanding of S100A4 function in different cell types in the PMN, single-cell RNA sequencing analysis was performed with the lungs from 4T1 tumor-bearing mice treated with IgG control or S100A4-11 antibody. The treatment scheme is identical to that shown in Fig. 3b. Dissociated single cells were pooled from the lungs of three mice per treatment group, to account for potential individual variation. 11,512 and 9,955 cells were captured from isotype control and the S100A4-11 antibody-treated lungs, respectively. UMAP clustering identified 25 cell clusters. Known lineage markers from the literature and single-cell atlases (**Figure 5a**, **Supplementary Figure 7a**-7b) (*45*) were used to identify clusters of endothelial cells, epithelial cells, granulocytes, T cells, B cells, NK cells, macrophages, and fibroblasts (**Figure 5a).** Comparison of the different cell-type proportions revealed that S100A4-11 treatment resulted in increased B cells, NK cells, and monocyte/macrophages in the lung while the proportions of T cells and granulocyte were unaffected (**Figure 5b, Supplementary Figure 7c**).

**Fig 5.**
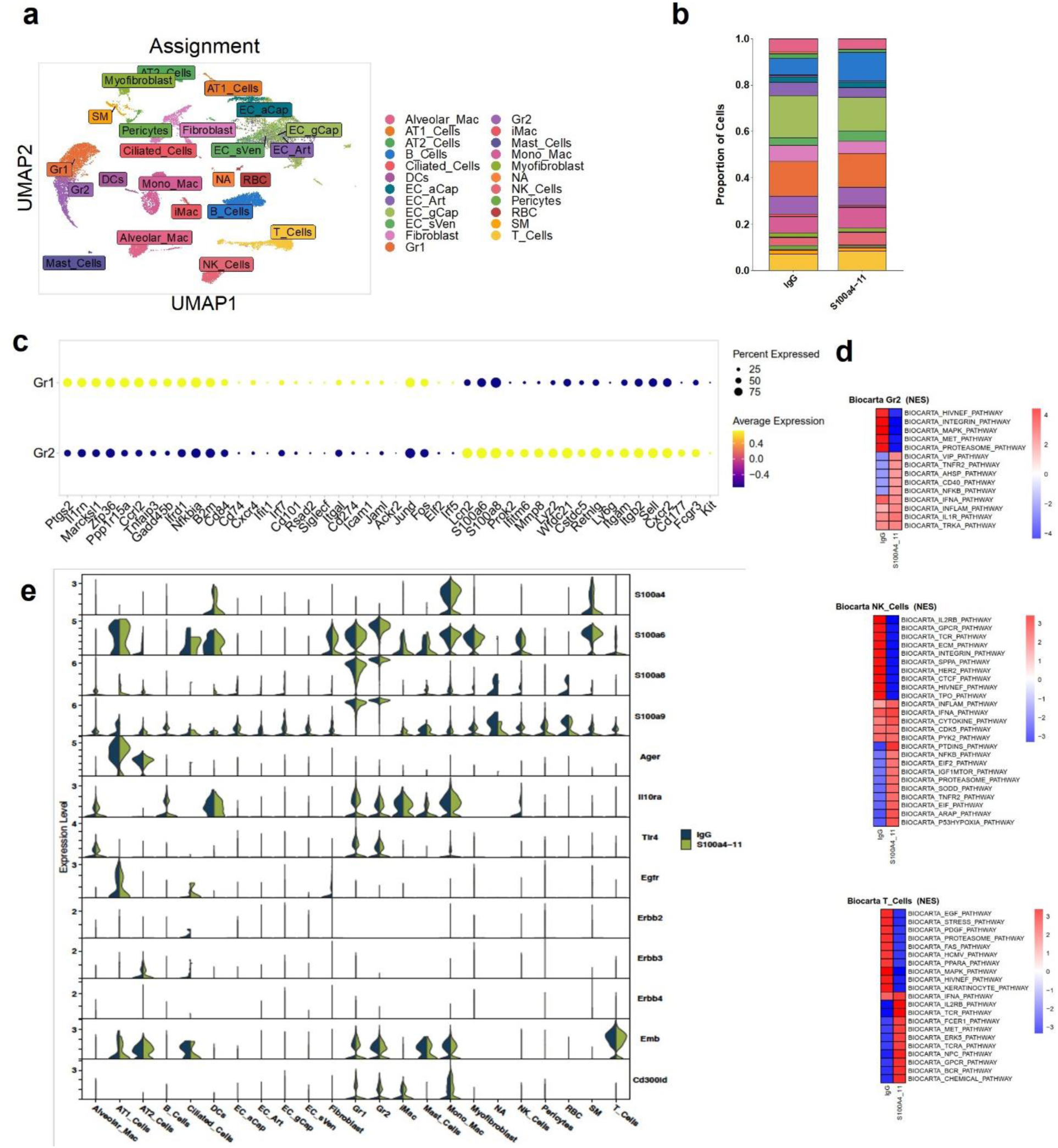
Single-cell RNA-seq analysis of day 14 lungs shows changes in multiple immune and stromal cell types upon S100A4-11 antibody treatment. (a) Single-cell RNA sequencing analysis was on the lung samples from 4T1 tumor-bearing mice after IgG (11,512 cells) and 4-11 (9,955 cells) antibody treatment. UMAP projection of all cells, color coded by assignment. (b) Bar graphs showing proportions of different cell types by sample. (c) Top differentially expressed genes and selected neutrophil marker gene expression in Gr1 and Gr2 granulocytes. (d) GSEA pathway analysis of differentially expressed genes in granulocytes (Gr2), NK cells and T cells, comparing IgG vs S100A4-11 antibody-treated mice. (e) Violin plot showing expression patterns of S100A family proteins (ex: S100A4/6/8/9) and their binding receptors.

Neutrophils/granulocytes formed clusters (C02 and C05) away from monocytes and macrophages (**Figure 5a**, **Supplementary Figure 7a).** Notably, there were two molecularly distinguishable subtypes of neutrophils/granulocytes (Gr1 and Gr2) in the lung (**Figure 6a, 6c**). The expression of classical markers of neutrophils in mice, *Ly6g*, *Cxcr2*, *S100a8, S100a9, Lcn2, Mmp8, CD177, Fcgr3* (*CD16*)*, Sell* (*CD62L*)*, Itgam/Cd11b*, was higher in Gr2 than in Gr1 cells. However, the expression of *Ptgs2/COX2, Cd84, Cxcr4* and *Nfkbia* was higher in Gr1 cells (**Figure 5c, Supplementary Figure 7d**). Gr1 cells also expressed higher levels of transcriptional factors (*JunD*, *Fos, Elf2, Irf5, and Irf7*) associated with pro-inflammatory neutrophils in the lung (*46, 47*). Moreover, the cells within Gr2 cluster expressed higher levels of markers associated with circulating neutrophils (Fcgr3, CD16^hi^, CXCR2^hi^, CXCR4^lo^, Sell (CD62L)^hi^) (*48–50*). The expression of *Ly6g* in Gr2 but not Gr1 combined with significantly reduced Ly6G^+^ granulocytes in the PMN (**Figure 3**) indicates that Gr2 neutrophil infiltration is blocked by S100A4-11 antibody treatment. Gene set enrichment analysis (GSEA) of matching cell types (Gr2, NK cells, and T cells) from the PMN of IgG vs. S100A4-11 treated lungs showed multiple differences in significantly enriched inflammation pathways, including tumor necrosis factor receptor 2 (TNFR2) and nuclear factor-kappa B (NF-ΚB), as well as the CD40 pathway (NES >2, **Figure 5d**) (*51*) in the Gr2 cells from S100A4-11 antibody-treated mice. NK cells from these mice also exhibited differential positive enrichment of inflammatory pathways (TNFR2 and NFKB) and T cells also showed enrichment of pathways associated with T cell activation (e.g., T-cell receptor (TCR) and IL-2 receptor B (IL2RB) pathways) (**Figure 5d**).

**Fig 6.**
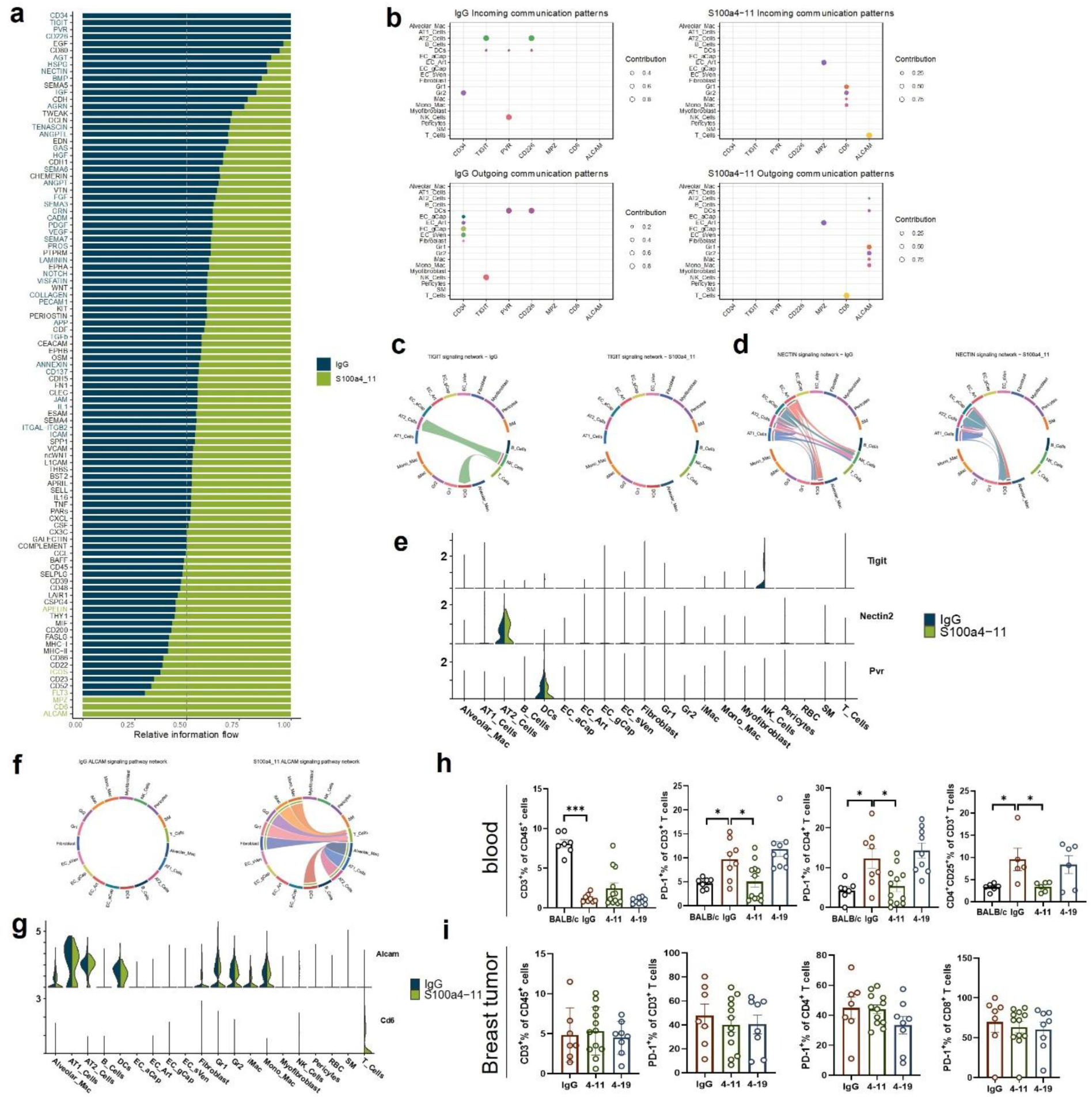
CellChat analysis of day 14 lungs show changes in multiple immune and stromal cell types upon S100A4-11 treatment. (a) Pan cell type information flow filtered for significantly enriched pathways in either sample, as determined by CellChat. Blue: enriched in IgG treated group; Green: enriched in 4-11 treated group. (b) Top differentially enriched pathways in either sample are selected and shown to depict differences in relative contributions of various cell types to outgoing and incoming signaling from each sample. (c, d) Chord diagram showing TIGIT-NECTIN signaling network strengths between IgG vs 4-11 treated group. (e) Violin plot showing expression levels of *TIGIT, NECTIN2, and PVR* in day 14 lung cells post-injection. Blue: IgG treated group; Green: S100A4-11 treated group. (f) Chord diagram showing CD6-ALCAM signaling network strengths between IgG vs 4-11 treated lung. (g) Violin plot showing expression levels of *Cd6 and Alcam*. (h) Flow cytometry analysis of blood showing all T cells (CD3^+^/CD45^+^), PD-1^+^/CD3^+^, PD-1^+^/CD4^+^ (BALB/c: n=7; tumor-bearing group: n=8-12), and CD4^+^CD25^+^/CD3^+^ T cells (n=5-6). Compared with IgG, S100A4-11 treatment significantly decreased PD-1^+^ cell fractions in both the CD3^+^ and CD4^+^ as well as the percentage of CD4^+^CD25^+^ T cells from the blood. (i) Flow cytometry analysis of breast tumors using the same gating strategy for T cells shown in (h) (n=7-12 per group). Ordinary one-way ANOVA analysis was used. Error bar denotes mean ± SEM. *\*p<*0.05*; ***p<*0.001.

To determine whether these changes were due to direct inhibition of S100A4 binding to its putative receptors, we analyzed the expression patterns of the S100A4/6/8/9 family members and known S100A4 binding receptors in the various lung cells (**Figure 5e**). Varied expression patterns of the S100A4 receptors, advanced glycosylation end product-specific receptor (AGER/RAGE), Toll-like receptor 4 (TLR4), IL-10 receptor subunit alpha (IL10RA), epidermal growth factor receptor (EGFR), Erb-B2 receptor tyrosine kinase 2 (ERBB2), ERBB3, embigin (EMB), and CD300 molecule-like family member D (CD300LD), were detected in a subset of lung epithelial cells, myeloid cells, and T cells. Notably, no significant differences in S100A4 receptor expression were detected in Gr1 and Gr2 granulocytes and T cells between the IgG control and S100A4-11 antibody-treated mice. However, reduced *Il10ra* and *S100a8* expression was detected in the NK cells of the S100A4-11 antibody-treated mice **(Figure 5e**).

### S100A4-11 treatment altered cell:cell signaling to and from T, NK, and neutrophils

Next, we explored how S100A4-11 antibody treatment and reduced neutrophil infiltration affect cell:cell communication among various cell types in the PMN. We used the CellChat algorithm (*31*) to analyze our single-cell RNA-seq data (**Figure 6**). Using known ligand-receptor interactions, CellChat predicts putative communication networks between two cell types. The CellChat analysis revealed overlapping and unique patterns of cell:cell communication between IgG vs S100A4-11 treated lung cells. There were significantly enriched signaling pathways that were present in the isotype control (IgG) but not in S1004-11 antibody-treated mice and vice versa (**Figure 6a**). When the top activated or repressed pathways in the lungs of the S100A4-11 antibody-treated mice were plotted, clear differences were observed in the signaling cell types and pathways in the S100A4-11 antibody-treated mice (**Figure 6b)**. Specifically, in the lung of the control IgG antibody-treated mice, CD34 signaling from endothelial cells to Gr2, TIGIT signaling from NK cells to AT2 and DCs, PVR from DCs to NK cells and CD226 signaling from DCs to AT2 cells signaling are present. However, these signaling patterns are absent in S100A4-11 antibody-treated mice (**Figure 6b-6e, Supplementary Figure 8a**-c). Moreover, CD6 signaling from T cells to myeloid cells and activated leukocyte cell adhesion molecule (ALCAM) signaling from myeloid cells to T cells were present in the lungs of the S100A4-11 antibody-treated mice but absent in the IgG antibody-treated mice. These results suggest that NK and T cells, which have known tumoricidal functions to suppress metastasis (*52, 53*), are reactivated in the lungs of the S100A4-11 antibody-treated mice.

Previous studies have reported that NK and T cells play a critical role in eliminating metastatic cancer cells in the lung (*54, 55*). CD6/ALCAM bi-directional signaling that amplifies T cell activation (*56, 57*), and this signaling is present in S100A4-11 treated but not in IgG treated mice (**Figure 6f, Supplementary Figure 8c**). Specifically, T cells in the lung of S100A4-11 treated mice expressed CD6, whereas this was not the case of the IgG treated mice (**Figure 6g**). In contrast, ALCAM was expressed in similar levels in alveolar epithelial cells, macrophages, DCs, fibroblasts, Gr1/Gr2 granulocytes and monocytes/macrophages from the mice of both treatment groups (**Figure 6g**), suggesting that S100A4-11 treatment activates the T cells. in addition, our CellChat analysis showed that TIGIT-Nectin interaction is altered in the S100A4-11 antibody-treated mice (**Figure 6c-6d**). TIGIT is an established NK and T-cell checkpoint molecule and is highly expressed in NK cells while TIGIT receptors PVR and NECTIN2 are expressed on AT2 and DCs in the IgG treated mice (**Figure. 6e).** However, TIGIT expression in NK cells is reduced in S100A4-11 treated lungs (**Figure 6e**), suggesting reactivation of NK cells in the PMN.

Together, our CellChat analyses are consistent with GSEA pathway analysis showing that T cells and NK cells are de-suppressed/activated upon S100A4-11 antibody treatment (**Figure. 6c-6d**). Difference in the signaling to and from fibroblasts, endothelial cells, dendric cells, and monocyte/macrophages were also observed (**Supplementary Figure 8d**-f), indicating that S100A4-11 treatment, directly or indirectly, affects critical signaling pathways among multiple cell types in the PMNs that promote immune suppression.

To further evaluate T cell activation in S100A4-11 treated mice, T cells from the blood of the mice were analyzed by flow cytometry. Compared to BALB/cJ control mice, all tumor-bearing mice had lymphopenia, indicated by significantly lower percentages of circulating CD3^+^ T cells (**Figure 6h**). However, the fraction of PD1^+^ T cells was significantly increased in the tumor-bearing mice (IgG control, **Figure 6h**). Following S100A4-11 treatment, circulating PD1^+^/CD3^+^ T cell frequency, including Tregs (CD4^+^CD25^+^), was significantly decreased, reaching frequencies similar to non-tumor-bearing mice (**Figure 6h, Supplementary Figure 8g**). There was no significant difference in the fraction of PD1^+^ cells among breast tumor associated T cells and circulating CD8^+^ T cells in any treatment group (**Figure 6i, Supplementary Figure 8h**). These results suggest that S100A4-11 treatment reduces the frequency of immune-suppressive T cells in the blood.

### Combined S100A4-11 and TIGIT inhibition blocks late lung metastasis

To determine the potential impact of S100A4-11 treatment in clinical settings, we tested whether S100A4-11 treatment can suppress breast metastasis during late stages of disease progression when the PMN is already established. To do this, we initiated the treatment with S100A4-11 in 4T1 mammary tumor bearing mice on day 14 after transplantation, when the PMN had established (**Figure 7a**). Although there was reduced number of lung nodules on average, S100a4-11 treatment alone did not reach statistically significant suppression of lung metastasis (**Figure 7b**). Similarly, anti-TIGIT monotherapy was not sufficient to significantly reduce lung metastasis (**Figure 7b**). However, a combination treatment using both S100A4-11 and TIGIT antibodies significantly reduced the number of lung nodules, even when the treatment was initiated post-PMN formation (**Figure 7b–7d**).

**Fig 7.**
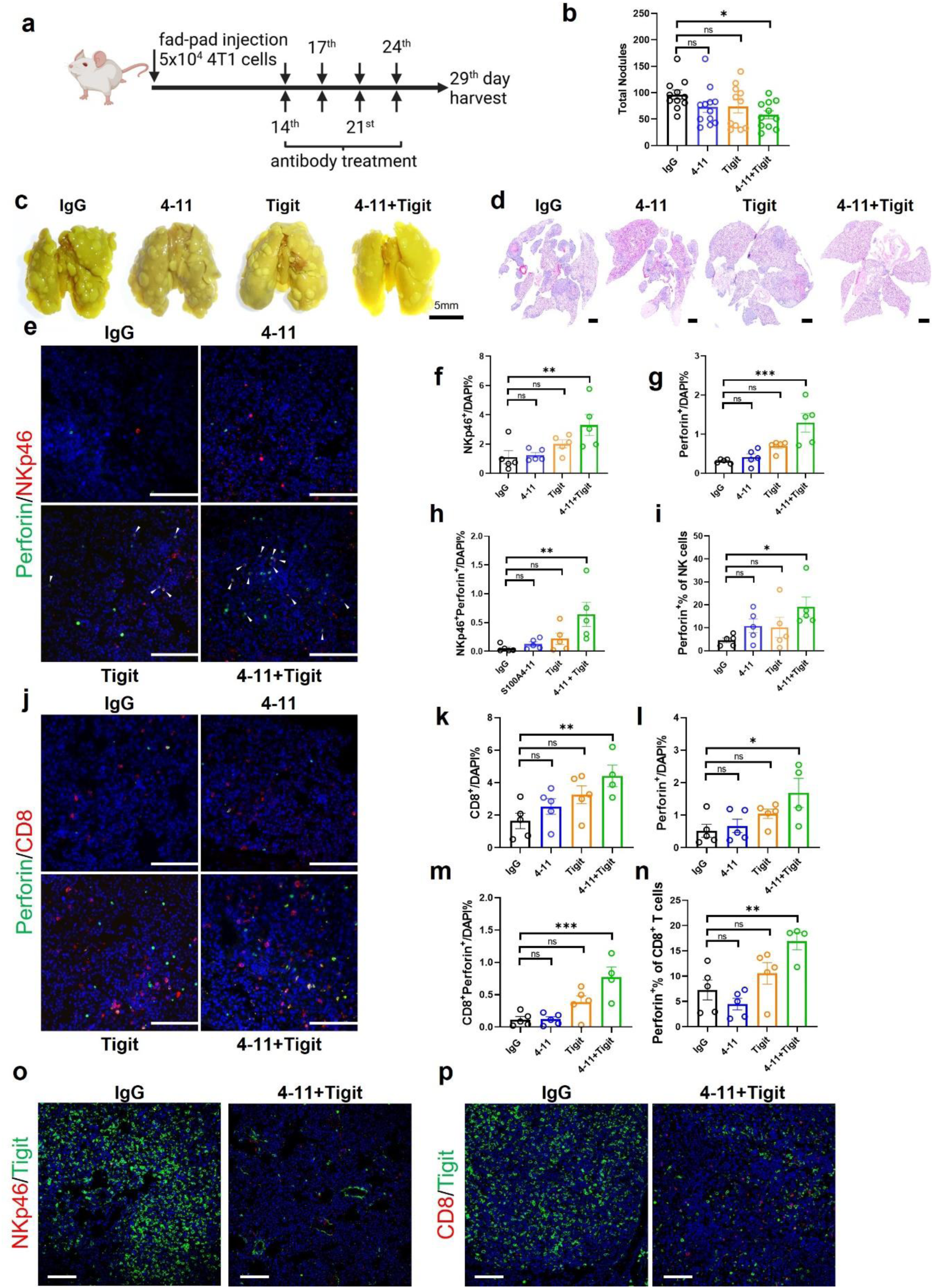
S100A4-11 plus Tigit antibody treatment suppresses late lung metastasis in the 4T1 breast cancer model. (a) S100A4-11 and Tigit combination antibody treatment was started on day 14 post-injection in 4T1 tumor-bearing mice. The mice were treated twice per week for 2 week. (b) Quantification of metastatic lung nodules from IgG, 4-11, Tigit, 4-11+Tigit antibody-treated mice (n=11-12). (c) Representative images with metastatic nodules from IgG, 4-11, Tigit, 4-11+Tigit antibody-treated mice. Scale bar denotes 5 mm. (d) Hematoxylin and eosin-stained overall lung tissues scanned by an Olympus VS200 (n=3). Scale bar indicates 1 mm. (e) Immunofluorescence analysis of NKp46/Perforin staining in lung tissue from 4T1 tumor-bearing mice. Arrows point out double-positive cells. Scale bar denotes 100µm. (f) NKp46^+^ and (g) Perforin^+^ (h) NKp46^+^Perforin^+^ cells are significantly increased in the 4-11+Tigit treated lungs, compared with IgG control (n=5). (i) Perforin^+^ percentage in NK cells is significantly increased in the lung of 4-11+Tigit treated mice, compared with IgG control. (n=5) (j) Immunofluorescence analysis of CD8/Perforin staining in the lung tissue of 4T1-bearing mice. Scale bar denotes 100µm. (k) CD8^+^ and (l) Perforin^+^ (m) CD8^+^Perforin^+^ cells are significantly increased in the 4-11+Tigit treated lungs, compared with IgG control (n=4-5). (n) Perforin^+^ percentage in CD8^+^ T cells is significantly increased in the lung of 4-11+Tigit treated mice, compared with IgG control (n=4-5). Three representative fields per mouse lung were counted and averaged. Datapoints represent average per mouse per group. Ordinary one-way ANOVA analysis was used. Error bar means ± SEM. *\*p<*0.05; *\*\*p<*0.01; *\*\*\*p<*0.001. (o, p) Late-stage metastatic lung tissues were stained with NKp46/Tigit (o) or CD8/Tigit (p) (n=3-4). Scale bar denotes 100µm.

Next, the mechanism of the combination treatment was evaluated by analyzing the markers of NK and T cell activation by double-immunofluorescence analysis. The combined therapy significantly increased the abundance of NK cells and CD8+ cytotoxic T cells in the lung (**Figure 7e-7f, 7j-k**). The combination treatment significantly increased the number of perforin-expressing, activated NK cells (NKp46^+^/Perforin^+^) (**Figure 7e–7i**) and CD8^+^ T cells (CD8^+^/Perforin^+^) (**Figure 7j–7n**), compared with the IgG antibody-treated mice. Although there were general trends toward increased NK and CD8^+^ T-cell numbers in the monotherapy-treated groups, these increases did not reach statistical significance. Concurrently, the number of TIGIT^+^ cells was drastically reduced in the lung nodules of the combination S100A4-11/TIGIT antibody-treated mice compared with the IgG antibody-treated mice (**Figure 7o, p**). On the other hand, the combination treatment post PMN formation had no effect on the number of neutrophils/PMN-MDSCs as anticipated (**Supplementary** Figure 9). These results indicate that the novel combination of S100A4 and TIGIT antibodies increases cytotoxic NK and T cell numbers in lung nodules to suppress lung metastasis, even when the treatment is initiated post-PMN formation.

## Discussion

Recent approval of Pembrolizumab for metastatic breast cancer highlights the importance of the immune system in breast cancer progression and metastasis (*58, 59*). Here we present data that our novel S100A4 blocking antibody significantly reduces lung metastases in both the spontaneous and transplantable mouse models of breast cancer (*35, 60*). Furthermore, we provide compelling evidence that S100A4-blocking antibody prevents PMNs formation by inhibiting neutrophil infiltration which in turn results in activation of cytotoxic NK and T cells. Importantly, a novel combination therapy targeting both S100A4 and TIGIT shows significant reduction of lung metastases even when the treatment is initiated after PMN formation, greatly expanding the clinical utility of this novel combination. Together with the observation that high S100A4 expression portends poor survival and is strongly associated with metastasis in multiple human cancers, this study provides compelling preclinical evidence for further development of our S100A4-11 blocking antibody to treat metastatic cancers.

S100A4 is a small calcium binding protein that forms dimers or multimers that bind to various target proteins (*61*). S100A4 protein can be detected in the nucleus, cytoplasm, or extracellular space, indicating both intracellular and extracellular functions. Intracellularly, it binds with non-muscle myosin II1 (NMIIA) to influence migration and metastasis (*62, 63*). It also promotes self-renewal and tumorigenic potential of cancer stem cells (*16, 60*). In gliomas, it has been demonstrated to be an upstream regulator of the master regulators of mesenchymal transition that control the epithelial-mesenchymal transition process (*16*). In non-cancer cells, S100A4 promotes immune-suppressive phenotypes in T cells and myeloid cells (*15*) and its expression in cancer-associated fibroblasts (CAF) is associated with metastasis (*64*). Extracellularly, S100A4 is released from cancer and non-cancer cells to promote angiogenesis and can function as a chemokine for immune cells (*65, 66*). In the TME, S100A4-induced proteins alter tumor cell mobility and regulates vasculature and extracellular matrix to promote tumor cell extravasation (*65, 67*). Thus, accumulating evidence indicates that S100A4 is not only involved in cancer stemness and cancer metastasis but mediates immune suppression in the TME.

Previous genetic and blocking antibody studies reported that S100A4 inhibition suppresses breast cancer metastasis to the lung (*19, 21, 22*). For example, a previous study reported that a blocking antibody against the mouse S100A4 protein, clone 6B2, inhibits lung metastasis in the MMTV-PyMT model by increasing T cell infiltration to the primary tumor and lung nodules (*21*). Unlike this study, we did not detect significant alterations in T-cell infiltration in the primary tumors or lung metastases of the S100A4-11 antibody-treated mice. Instead, our S100A4-11 antibody significantly blocks neutrophil infiltration in pre-metastatic lungs (**Figure 2e**), thereby activating T and NK cells in the PMN. This presumably results in more efficient elimination of infiltrating cancer cells (**Figure 6b-6f**, **Figure 7e-7n**). In the primary tumor, the only significant alteration we observed from S100A4-11 treatment is in the frequency of CD163^+^, but not CD206^+^, macrophages (**Figure 1m**, **Figure 2h**). Our results suggest that the major effect of S100A4-11 treatment is mediated through a paracrine effect that modulates the infiltration of neutrophils that are necessary for PMN formation in the lung.

Mechanistically, we discovered that S100A4-11 treatment affects multiple immune and stromal cell types in the primary tumor, lung, and blood at different stages of disease progression. The importance of neutrophils, MDSCs, NK cells and T cells in promoting metastasis and PMN formation is well established (*68*). A high neutrophil-to-lymphocyte ratio indicates poor survival in multiple cancer types (*69*). Importantly, neutrophils retain T and NK cells in an immunosuppressive state in the PMN (*70*). Notably, the most drastic phenotype we observe in pre-metastatic lung tissue following anti-S1004-11 antibody treatment is a significantly reduced number of infiltrating neutrophils and the subsequent activation of T cells via CD6/ALCAM signaling and NK cells via the loss of TIGIT signaling, as determined by single cell RNA-sequencing and CellChat analysis (**Figure 6b–6f**). Both our proteomics (**Figure 4, Supplementary** Figure 6) and single-cell RNA-sequencing analyses (**Figure 5**) support the conclusion that anti-S100A4-11 antibody treatment reduces the number and function of lung infiltrating neutrophils (Ly6G^+^) in the PMN.

Others have reported that PMN-MDSCs/neutrophils progressively accumulate in the lung of 4T1 tumor-bearing mice (*71*) and provide a carbon source for tumor growth (*3*). The PMN is established by bone marrow derived PMN-MDSCs/neutrophils before tumor cells infiltrate the lung (*45, 72*). Here, no significant alterations in PMN-MDSC/neutrophil frequency are observed in the primary tumor or blood in 4T1 model. However, a specific subset (Gr2) of granulocytes (Ly6G^+^/S100A8/9^+^) is reduced by S100A4-11 antibody treatment (**Figure 3l, 3o–3p**). An earlier study suggested a positive feedback loop through TLR4 dependent pathway for PMN formation (*38*), and a different study showed that S100A4 promotes MDSC survival through TLR4 signaling (*7*). Hence, S100A4 blocking antibody may interfere with S100A4/TLR4 signaling axis, which may be important for granulocytic MDSC survival and pre-metastatic niche formation. However, examination of known S100A4 receptors has revealed that Gr1 (unaffected) and Gr2 (affected) cells have similar *Tlr4, Emb*, and *Il10ra* RNA expression levels. Based on marker expression and cell number changes, we conclude that Gr1 represents lung-resident neutrophils, whereas Gr2 represents neutrophils that are recruited to the lungs by breast tumor cells to prime the lungs for metastasis in a S100A4 dependent manner (directly or indirectly).

Our single cell analysis revealed that S100A4-11 treatment may suppress lung metastasis through indirectly activating T cells and NK cells that have been demonstrated to kill metastatic cancer cells (*54, 73*). Therefore, we propose that blocking the soluble S100A4 function at early stages of breast cancer can reduce metastatic burden in the lung by 1) reducing neutrophil infiltration to prevent successful PMN formation and 2) reprogramming neutrophils and other cell types in the lung, directly or indirectly, that lead to de-repressed cytotoxic NK cells and T cells that then are able to eliminate metastatic cancer cells. At later stages (post PMN formation), S100A4-11 treatment is insufficient to change the neutrophil number in the lung; however, when combined with anti-TIGIT antibody treatment, it significantly reduces the number of lung nodules.

This study also revealed a previously unknown TIGIT signaling between endothelial cells and NK cells in the lung. Consistently, adding anti-TIGIT treatment to S100A4-11 treatment in late stages of metastases increases T and NK cell numbers and expression of a cytotoxic molecule perforin in the lung tissue, resulting in significantly reduced lung metastases. However, anti-TIGIT treatment is not sufficient to reduce lung metastasis on its own after PMNs have been established. We propose that a novel combination of S100A4 and TIGIT inhibition reprograms the TME to enable full functionality of cytotoxic NK and T cells to eliminate nearby cancer cells (micro metastases) and prevent macro-metastasis growth.

S100A4-11 treatment was well tolerated in two distinct mouse models of breast cancer in two different genetic backgrounds (**Supplementary Figure 3g, 3i**). In addition, *S100a4^-/-^* mice are viable and fertile (*19, 74*), suggesting that S100A4-11 is likely to have very low toxic side effects in the clinic. The S100A4-11 clone has approximately 10-fold higher affinity for the human S100A4 protein than mouse protein (**Supplementary Figure 2f**-g), strongly supporting its anticipated efficacy in humans.

In summary, this study provides evidence for a novel mechanism of S100A4 in PMN formation, complex signaling patterns among different cell types in the premetastatic lung, and the therapeutic potential of targeting S100A4 as a novel immunotherapy to suppress cancer metastasis in early and late stages of breast cancer progression. Additionally, in the clinical setting, breast cancer patients are often diagnosed with late-stage disease where circulating tumor cells are detectable in the blood and the PMN is already established (*75*). The combined S100A4-11 and anti-TIGIT treatment significantly reduces lung metastases even when the treatment is initiated post-PMN formation and dramatically reactivates cytotoxic NK cells and CD8^+^ T cells even in late metastasis stage. Overall, this study provides a novel insight into the role of S100A4 in breast cancer metastases at different stages of disease and promising preclinical results from targeting S100A4 as a monotherapy or a combination therapy with TIGIT.

## Materials and Methods

### Mice

All animal studies were conducted in accordance with the protocol approved by the Institutional Animal Care and Use Committee (IACUC) at the Houston Methodist Research institute (Protocol #AUP=0120-0003). BALB/cJ (JAX#000651) and MMTV-PyMT (JAX#002374) mice were purchased from the Jackson Laboratory. *S100a4^-/-^* (FSP1^GFP/GFP^) mice (*27*) were purchased from JAX (#012904). MMTV-PyMT spontaneous breast tumor model in FVB background were purchased from JAX (#001800). All mice were housed and cared for by the staff in the Houston Methodist Research Institute Comparative Medicine Program.

### Cell lines

4T1 and E0771 breast cancer cell lines, respectively derived from breast tumors in the BALB/c and C57BL/6 genetic backgrounds, respectively, were purchased from American Type Culture Collection (ATCC) and cultured in 10% fetal bovine serum (FBS)/Roswell Park Memorial Institute (RPMI) 1640 medium or Dulbecco’s Modified Eagle Medium (DMEM) with penicillin/streptomycin (GenDepot, Cat#CA005-010).

### Tumor models and in vivo treatments

For 4T1 allografts using BALB/c females, 5×10^4^ cells in 50 µl of phosphate-buffered saline (PBS) were injected into the 2^nd^ right mammary fat-pad. Isotype IgG control or anti-S100A4 blocking antibody were injected (10mg/kg, IP, one time per week) starting on 3^rd^ day post-injection. 4T1 mammary tumor burden was measured twice weekly, using the formula ½ x (length x width^2^). For PMN analysis, mice were euthanized on day 14 after injection and lungs were harvested. For end point analyses, all mice were harvested on day 29.

To test anti-S100A4 antibodies in a spontaneous breast tumor model, 6-week-old MMTV-PyMT female mice were treated with IgG or anti-S100A4 antibody clones via tail-vein injection.

Mice were treated with isotype control or anti-S100A4 blocking antibody clones (10mg/kg, twice a week for 4 weeks). All mice were euthanized when primary tumors reached humane endpoints at approximately 12∼14 weeks of age.

For E0771 allografts using female C57BL/6 or *Fsp1-*knockin heterozygous and homozygous mice (purchased from JAX and maintained by crossing with C57BL/6 mice), 5 × 10^5^ cells in 50 μL of PBS were injected into the second right mammary fat pad. The E0771 mammary tumor burden was measured twice a week, and tumors were harvested on day 21 post-injection.

### Chick neurite outgrowth Assay

Briefly, isolated chick dorsal root ganglia (DRG) were cultured in collagen, supplemented with recombinant mouse S100A4 protein (25 µg/ml) with or without S100A4 antibodies. After incubation for 3 days, the DRG were fixed and stained with ß-3-tubulin and neurite lengths were measured and compared with those of the control (**Supplementary** Figure. 2).

### Immunohistochemistry and Immunofluorescence analysis

Breast tumor and lung tissues were harvested and fixed in 10% formalin solution and embedded in paraffin. The samples were sectioned at a 5 µm thickness, deparaffined, and boiled in 10 mM sodium citrate (pH6.0) for retrieve antigens. The sections were blocked with 5% goat serum for 20-30 min and incubated with primary antibody overnight at 4°C (**Supplementary Table 1**). After washing, the sections were incubated with a biotinylated secondary antibody for 45-60 min, followed by Avidin-biotin complex (ABC) solution (Vector Laboratory, Cat# PK-4000) for 1 hour. The 3,3′-diaminobenzidine (DAB) chromogenic reaction was used for visualization. Nuclei were counterstained with hematoxylin. Images were taken by Zeiss Primostar 3 microscope. For immunofluorescence analysis, Alex fluor 488/555/594 secondary antibodies were used, and nuclei were stained with 4′,6-diamidion-2-phenylindole (DAPI). To minimize background autofluorescence, the autofluorescence quenching kit (Vector Laboratory, Cat# SP-8500) was applied. Zeiss Axiovert 200M invert fluorescence microscope, Leica DM6 microscope, and FV3000 confocal microscopy (Olympus) were used for imaging.

### Flow cytometry

Mouse breast tumor and lung tissues were dissected, cut into small pieces, and incubated in Accutase/Collagenase/Hyaluronidase enzyme mixture for 15-30 min at 37°C. After removing the enzymes, tissue was resuspended in PBS for mechanical dissociation by pipetting to generate a single-cell suspension. Blood was also harvested from the same mice and washed to make single-cell suspension of peripheral blood mononuclear cells (PBMCs). Before staining, red blood cells (RBCs) were removed in RBC lysis buffer (Biolegend), and the cells were filtered through a 40µm cell strainer. Cells were first incubated with anti-mouse CD16/32 blocking antibody (Biolegend cat#156604) for 5 minutes on ice and then stained with validated antibodies (**Supplementary Table 2**) following a standard protocol. A cell viability dye (Cat#65-0863-14, Invitrogen) was used to distinguish live/dead cells. All stained samples were run through BD LSRII flow cytometry using the same instrument settings and were analyzed by Flowjo v10.9.0.

### Quantitative PCR

Tissues were lysed in TRIzol (ThermoFisher), RNA was extracted and reverse transcribed into complementary DNA (cDNA) using the iScript^TM^ Reverse Transcription Supermix (Bio-Rad). Quantitative (qPCR) was performed using CFX96 PCR detection system (Bio-Rad) and qPCR master mix (GenDepot), according to the manufacturer’s instruction. The following primers were used: 18s-FW 5’ GAGGGAGCCTGAGAAACGG 3’; 18s-RV 5’ GTCGGGAGTGGGTAATTTGC 3’; S100a4-FW 5’ GTCCACCTTCCACAAATACTC 3’; S100a4-RV 5’ GCAGCTTCATCTGTCCTTT 3’; S100a8-FW 5’ CTGAGTGTCCTCAGTTTGTG 3’; S100a8-RV 5’ CCTTGTGGCTGTCTTTGT 3’; S100a9-FW 5’ ATGGTGGAAGCACAGTTG 3’; S100a9-RV 5’ AGCTGATTGTCCTGGTTTG 3’; Mmp9-FW 5’ CAACAGCTGACTACGATAAGG 3’; Mmp9-RV 5’ CTCAAAGATGAACGGGAACA 3’.

### Single-cell RNA-sequencing analysis

For single-cell RNA-sequencing, lungs were isolated at day 14 from BALB/cJ female mice bearing 4T1 tumors in their mammary fat pads and treated with either IgG or anti-S100A4 antibody. The lungs were then dissociated. Single cells from three mice per treatment group were pooled together before capture on a 10× Chromium instrument, and single-cell sequencing libraries were generated using Chromium Next GEM Single Cell 5′ Reagent Kit v2 (Cat#PN-1000263). The data were analyzed using the established pipeline. Briefly, CellRanger 7.1.0 (RRID: SCR_022697) was used to align the reads, low-quality single cells were identified by miQC (RRID:SCR_022697) with a posterior cutoff value of 0.8, and doublets were removed with scDblFinder (RRID:SCR_022700). Following the removal of low-quality and doublet cells, single cells were normalized and clustered using Seurat V4.3.0.1 (RRID:SCR_016341) and batch-corrected using Harmony v1.0 (RRID:SCR_022206). Subsequently, the dimensionality of the data was reduced using principal component analysis based on the 2,000 most variably expressed genes. Batch-corrected principal components were used as inputs for Louvain-based graphing with a resolution of 0.1. Cluster-specific gene expression was then used for cell-type annotation. For endothelial cell clusters, the Azimuth-Human Lung v2 reference (RRID:SCR_021084) was used to auto-assign endothelial cell subtypes. Any auto-assigned subtypes with less than 100 cells were removed.

### Cell-to-cell communication analysis using CellChat

CellChat (RRID:SCR_021946) analysis was used to identify cell-to-cell interactions across different cell types. Cell types with less than 100 cells from the CellChat analysis were excluded, and the ligands and receptors that were expressed in at least 10% of the population were considered (default cutoff). CellChat then computed the probability of communication at the signaling pathway level with a threshold *p*-value of 0.05 for determining significant interactions. Finally, using the CellChat visualization tool plot, significant interactions were identified through chord plots, heatmaps, and bubble plots. Violin plots of ligand-receptor pairs were plotted based on Seurat.

### Statistical analysis

To compare the mean/average number of two independent treated groups or clusters, an unpaired *S*tudent’s *t-*test was applied, using GraphPad Prism software. Data is shown along with the mean ± standard error of the mean (SEM). One-way ANOVA analysis was used for multiple-group analysis.

## Supplementary Materials and Methods

**Expression and purification of S100A4 mAbs**

**Binding affinity of anti-S100A4 mAb using Bio-layer interferometry (BLI) Octet instrument**

**Measurement of anti-S100A4 mAb binding using ELISA**

**Chick DRG neurite growth assay**

**Pharmacokinetic evaluation of S100A4 antibody**

### Supplementary Figures

**Supplementary Fig 1. S100A4^+^/GFP^+^ expression comparison between blood and tumor stromal cells in the E0771 tumor-bearing *S100a4*^+/-^ mice**

**Supplementary Fig 2. Anti-human S100A4 antibody screen design and characterization**

**Supplementary Figure 3. Pharmacokinetic profile and tolerance of S100A4-11 antibody**

**Supplementary Figure 4. Effects of S100A4-11 antibody treatment on the MDSCs in the MMTV-PyMT primary breast tumor**

**Supplementary Figure 5. Effects of S100A4-11 antibody treatment on the blood and tumor MDSCs in 4T1 breast cancer model**

**Supplementary Fig 6. Cytokine array analysis of lung lysates from 4T1 tumor-bearing mice**

**Supplementary Figure 7. Cell type assignments of single-cell RNA-sequencing clusters**

**Supplementary Figure 8. Differential cell:cell communications among various cell types in the pre-metastatic lung**

**Supplementary Figure 9. S100A4-11 plus Tigit antibody treatment has no effect on the number of infiltrating neutrophils in late-stage metastatic lung**

### Supplementary Tables

**Supplementary Table1: The antibodies list used for IHC/IF and western blotting**

**Supplementary Table2: The antibodies list used for flow cytometry References (76–78)**

## Acknowledgement

We thank David Haviland and Matthew Vasquez at Houston Methodist Research Institute (HMRI) flow cytometry and Microscopy core facilities for their assistance. We also thank the staff in the HMRI Comparative Medicine Program for animal care and assistance.

## Funding

This work was supported partly by NIH R015R01NS121405-02 and 1R01CA271682-01A1 (KY), Golfers Against Cancer Foundation (KY), and CPRIT Core Grants RP150551 and RP190561 (ZA) and the Welch Foundation (AU-0042-20030616 to ZA).

## Contributions

JSL conducted all animal studies and prepared manuscript, HD conducted antibody cloning and assay evaluations; XW carried out antibody expression, purification, and characterization; HNT generated and analyzed single cell RNA-sequencing data, TW, NA, JBS, JAM, RK, CH, FL assisted with data collection and analyses, CDC and HKL provided chick embryos, ZA and NZ supervised antibody design and characterization and reviewed manuscript. KY conceived the study, supervised experimental design and data analyses, conducted antibody screens, and prepared the manuscript. All authors reviewed the manuscript.

## Conflict of interest statement

KY is a cofounder of EMPIRI, inc. HD, WX, TW, ZA, NZ, and KY are inventors of a patent application based on this work.

## References

1. H. Sung et al., Global Cancer Statistics 2020: GLOBOCAN Estimates of Incidence and Mortality Worldwide for 36 Cancers in 185 Countries. CA Cancer J Clin 71, 209–249 (2021).

2. K. Ganesh, J. Massague, Targeting metastatic cancer. Nat Med 27, 34–44 (2021).

3. J. Fares, M. Y. Fares, H. H. Khachfe, H. A. Salhab, Y. Fares, Molecular principles of metastasis: a hallmark of cancer revisited. Signal Transduct Target Ther 5, 28 (2020).

4. S. Paget, The distribution of secondary growths in cancer of the breast. 1889. Cancer Metastasis Rev 8, 98–101 (1989).

5. F. Veglia, E. Sanseviero, D. I. Gabrilovich, Myeloid-derived suppressor cells in the era of increasing myeloid cell diversity. Nat Rev Immunol 21, 485–498 (2021).

6. K. Li et al., Myeloid-derived suppressor cells as immunosuppressive regulators and therapeutic targets in cancer. Signal Transduct Target Ther 6, 362 (2021).

7. J. Albrengues et al., Neutrophil extracellular traps produced during inflammation awaken dormant cancer cells in mice. Science 361, (2018).

8. S. K. Wculek, I. Malanchi, Neutrophils support lung colonization of metastasis-initiating breast cancer cells. Nature 528, 413–417 (2015).

9. S. B. Coffelt et al., IL-17-producing gammadelta T cells and neutrophils conspire to promote breast cancer metastasis. Nature 522, 345–348 (2015).

10. Z. Granot et al., Tumor entrained neutrophils inhibit seeding in the premetastatic lung. Cancer Cell 20, 300–314 (2011).

11. M. Kowanetz et al., Granulocyte-colony stimulating factor promotes lung metastasis through mobilization of Ly6G+Ly6C+ granulocytes. Proc Natl Acad Sci U S A 107, 21248–21255 (2010).

12. Y. Ouyang et al., Tumor-associated neutrophils suppress CD8(+) T cell immunity in urothelial bladder carcinoma through the COX-2/PGE2/IDO1 Axis. Br J Cancer 130, 880–891 (2024).

13. A. Spiegel et al., Neutrophils Suppress Intraluminal NK Cell-Mediated Tumor Cell Clearance and Enhance Extravasation of Disseminated Carcinoma Cells. Cancer Discov 6, 630–649 (2016).

14. T. M. Ismail et al., S100A4 Elevation Empowers Expression of Metastasis Effector Molecules in Human Breast Cancer. Cancer Res 77, 780–789 (2017).

15. N. Abdelfattah et al., Single-cell analysis of human glioma and immune cells identifies S100A4 as an immunotherapy target. Nat Commun 13, 767 (2022).

16. K. H. Chow et al., S100A4 Is a Biomarker and Regulator of Glioma Stem Cells That Is Critical for Mesenchymal Transition in Glioblastoma. Cancer Res 77, 5360–5373 (2017).

17. Y. G. Cho et al., Overexpression of S100A4 is closely associated with progression of colorectal cancer. World J Gastroenterol 11, 4852–4856 (2005).

18. S. Gongoll et al., Prognostic significance of calcium-binding protein S100A4 in colorectal cancer. Gastroenterology 123, 1478–1484 (2002).

19. B. Grum-Schwensen et al., Lung metastasis fails in MMTV-PyMT oncomice lacking S100A4 due to a T-cell deficiency in primary tumors. Cancer Res 70, 936–947 (2010).

20. S. Liu et al., Anti-S100A4 antibody administration alleviates bronchial epithelial-mesenchymal transition in asthmatic mice. Open Med (Wars) 18, 20220622 (2023).

21. B. Grum-Schwensen et al., S100A4-neutralizing antibody suppresses spontaneous tumor progression, pre-metastatic niche formation and alters T-cell polarization balance. BMC Cancer 15, 44 (2015).

22. J. Klingelhofer et al., Anti-S100A4 antibody suppresses metastasis formation by blocking stroma cell invasion. Neoplasia 14, 1260–1268 (2012).

23. W. Tang, J. Chen, T. Ji, X. Cong, TIGIT, a novel immune checkpoint therapy for melanoma. Cell Death Dis 14, 466 (2023).

24. Q. Zhang et al., Blockade of the checkpoint receptor TIGIT prevents NK cell exhaustion and elicits potent anti-tumor immunity. Nat Immunol 19, 723–732 (2018).

25. L. H. Butterfield, Y. G. Najjar, Immunotherapy combination approaches: mechanisms, biomarkers and clinical observations. Nat Rev Immunol 24, 399–416 (2024).

26. K. E. Yost, H. Y. Chang, A. T. Satpathy, Recruiting T cells in cancer immunotherapy. Science 372, 130–131 (2021).

27. C. Xue, D. Plieth, C. Venkov, C. Xu, E. G. Neilson, The gatekeeper effect of epithelial-mesenchymal transition regulates the frequency of breast cancer metastasis. Cancer Res 63, 3386–3394 (2003).

28. Z. Fang, N. Forslund, K. Takenaga, E. Lukanidin, E. N. Kozlova, Sensory neurite outgrowth on white matter astrocytes is influenced by intracellular and extracellular S100A4 protein. J Neurosci Res 83, 619–626 (2006).

29. R. Donato, S100: a multigenic family of calcium-modulated proteins of the EF-hand type with intracellular and extracellular functional roles. Int J Biochem Cell Biol 33, 637–668 (2001).

30. E. Y. Lin et al., Progression to malignancy in the polyoma middle T oncoprotein mouse breast cancer model provides a reliable model for human diseases. Am J Pathol 163, 2113–2126 (2003).

31. S. Jin et al., Inference and analysis of cell-cell communication using CellChat. Nat Commun 12, 1088 (2021).

32. V. Bronte et al., Recommendations for myeloid-derived suppressor cell nomenclature and characterization standards. Nat Commun 7, 12150 (2016).

33. A. Mantovani, P. Allavena, F. Marchesi, C. Garlanda, Macrophages as tools and targets in cancer therapy. Nat Rev Drug Discov 21, 799–820 (2022).

34. A. K. Mehta, S. Kadel, M. G. Townsend, M. Oliwa, J. L. Guerriero, Macrophage Biology and Mechanisms of Immune Suppression in Breast Cancer. Front Immunol 12, 643771 (2021).

35. S. Yang, J. J. Zhang, X. Y. Huang, Mouse models for tumor metastasis. Methods Mol Biol 928, 221–228 (2012).

36. Y. Heng et al., CD206(+) tumor-associated macrophages interact with CD4(+) tumor-infiltrating lymphocytes and predict adverse patient outcome in human laryngeal squamous cell carcinoma. J Transl Med 21, 167 (2023).

37. S. M. Hacking et al., Microinvasive breast cancer and the role of sentinel lymph node biopsy. Sci Rep 12, 12391 (2022).

38. S. Hiratsuka et al., The S100A8-serum amyloid A3-TLR4 paracrine cascade establishes a pre-metastatic phase. Nat Cell Biol 10, 1349–1355 (2008).

39. Y. Wang et al., Glycolytic neutrophils accrued in the spleen compromise anti-tumour T cell immunity in breast cancer. Nat Metab 5, 1408–1422 (2023).

40. H. Lorchner et al., Neutrophils for Revascularization Require Activation of CCR6 and CCL20 by TNFalpha. Circ Res 133, 592–610 (2023).

41. Z. A. Lopez-Bujanda et al., Castration-mediated IL-8 promotes myeloid infiltration and prostate cancer progression. Nat Cancer 2, 803–818 (2021).

42. P. Gough, S. Ganesan, S. K. Datta, IL-20 Signaling in Activated Human Neutrophils Inhibits Neutrophil Migration and Function. J Immunol 198, 4373–4382 (2017).

43. M. Owyong et al., MMP9 modulates the metastatic cascade and immune landscape for breast cancer anti-metastatic therapy. Life Sci Alliance 2, (2019).

44. J. F. Lo et al., The epithelial-mesenchymal transition mediator S100A4 maintains cancer-initiating cells in head and neck cancers. Cancer Res 71, 1912–1923 (2011).

45. Y. Xiao et al., Cathepsin C promotes breast cancer lung metastasis by modulating neutrophil infiltration and neutrophil extracellular trap formation. Cancer Cell 39, 423–437 e427 (2021).

46. Z. Ai, Revealing key regulators of neutrophil function during inflammation by re-analysing single-cell RNA-seq. PLoS One 17, e0276460 (2022).

47. T. E. Khoyratty et al., Distinct transcription factor networks control neutrophil-driven inflammation. Nat Immunol 22, 1093–1106 (2021).

48. S. H. Bongers et al., Kinetics of Neutrophil Subsets in Acute, Subacute, and Chronic Inflammation. Front Immunol 12, 674079 (2021).

49. M. Bai et al., CD177 modulates human neutrophil migration through activation-mediated integrin and chemoreceptor regulation. Blood 130, 2092–2100 (2017).

50. K. J. Eash, J. M. Means, D. W. White, D. C. Link, CXCR4 is a key regulator of neutrophil release from the bone marrow under basal and stress granulopoiesis conditions. Blood 113, 4711–4719 (2009).

51. R. H. Vonderheide, Prospect of targeting the CD40 pathway for cancer therapy. Clin Cancer Res 13, 1083–1088 (2007).

52. M. A. Harris et al., Towards targeting the breast cancer immune microenvironment. Nat Rev Cancer 24, 554–577 (2024).

53. N. K. Wolf, D. U. Kissiov, D. H. Raulet, Roles of natural killer cells in immunity to cancer, and applications to immunotherapy. Nat Rev Immunol 23, 90–105 (2023).

54. I. S. Chan, A. J. Ewald, The changing role of natural killer cells in cancer metastasis. J Clin Invest 132, (2022).

55. K. Nakamura, M. J. Smyth, Immunoediting of cancer metastasis by NK cells. Nat Cancer 1, 670–671 (2020).

56. S. A. Chalmers et al., The CD6/ALCAM pathway promotes lupus nephritis via T cell-mediated responses. J Clin Invest 132, (2022).

57. A. W. Zimmerman et al., Long-term engagement of CD6 and ALCAM is essential for T-cell proliferation induced by dendritic cells. Blood 107, 3212–3220 (2006).

58. H. S. Rugo et al., Abemaciclib in combination with pembrolizumab for HR+, HER2-metastatic breast cancer: Phase 1b study. NPJ Breast Cancer 8, 118 (2022).

59. P. Schmid et al., Pembrolizumab for Early Triple-Negative Breast Cancer. N Engl J Med 382, 810–821 (2020).

60. G. Francia, W. Cruz-Munoz, S. Man, P. Xu, R. S. Kerbel, Mouse models of advanced spontaneous metastasis for experimental therapeutics. Nat Rev Cancer 11, 135–141 (2011).

61. F. Fei, J. Qu, M. Zhang, Y. Li, S. Zhang, S100A4 in cancer progression and metastasis: A systematic review. Oncotarget 8, 73219–73239 (2017).

62. B. Kiss et al., Crystal structure of the S100A4-nonmuscle myosin IIA tail fragment complex reveals an asymmetric target binding mechanism. Proc Natl Acad Sci U S A 109, 6048–6053 (2012).

63. E. J. Kim, D. M. Helfman, Characterization of the metastasis-associated protein, S100A4. Roles of calcium binding and dimerization in cellular localization and interaction with myosin. J Biol Chem 278, 30063–30073 (2003).

64. B. Schmidt-Hansen et al., Functional significance of metastasis-inducing S100A4(Mts1) in tumor-stroma interplay. J Biol Chem 279, 24498–24504 (2004).

65. J. T. O’Connell et al., VEGF-A and Tenascin-C produced by S100A4+ stromal cells are important for metastatic colonization. Proc Natl Acad Sci U S A 108, 16002–16007 (2011).

66. N. Ambartsumian et al., The metastasis-associated Mts1(S100A4) protein could act as an angiogenic factor. Oncogene 20, 4685–4695 (2001).

67. I. J. Bettum et al., Metastasis-associated protein S100A4 induces a network of inflammatory cytokines that activate stromal cells to acquire pro-tumorigenic properties. Cancer Lett 344, 28–39 (2014).

68. H. Peinado et al., Pre-metastatic niches: organ-specific homes for metastases. Nat Rev Cancer 17, 302–317 (2017).

69. A. J. Templeton et al., Prognostic role of neutrophil-to-lymphocyte ratio in solid tumors: a systematic review and meta-analysis. J Natl Cancer Inst 106, dju124 (2014).

70. Z. Gong, et al., Immunosuppressive reprogramming of neutrophils by lung mesenchymal cells promotes breast cancer metastasis. Sci Immunol 8, eadd5204 (2023).

71. M. Bosiljcic et al., Targeting myeloid-derived suppressor cells in combination with primary mammary tumor resection reduces metastatic growth in the lungs. Breast Cancer Res 21, 103 (2019).

72. S. Tabaries et al., Granulocytic immune infiltrates are essential for the efficient formation of breast cancer liver metastases. Breast Cancer Res 17, 45 (2015).

73. D. Krijgsman, M. Hokland, P. J. K. Kuppen, The Role of Natural Killer T Cells in Cancer-A Phenotypical and Functional Approach. Front Immunol 9, 367 (2018).

74. B. Grum-Schwensen et al., Suppression of tumor development and metastasis formation in mice lacking the S100A4(mts1) gene. Cancer Res 65, 3772–3780 (2005).

75. D. Lin et al., Circulating tumor cells: biology and clinical significance. Signal Transduct Target Ther 6, 404 (2021).

