## Supplementary Methods and Figures for "Combined inhibition of S100A4 and TIGIT suppresses late-stage breast cancer metastasis to the lung by activating T and NK cells"

#### Supplementary Materials and Methods

##### Expression and purification of S100A4 mAbs

Selected S100A4 binding hits were expressed as full-length IgG by fusion with rabbit or human constant regions in a mammalian expression vector using human embryonic kidney (HEK293) cells (Invitrogen) as we reported previously(*76, 77*) . Antibodies were purified with protein A affinity resin using a fast protein liquid chromatography system (FPLC, GE) (*76-78*).

##### Binding affinity of anti-S100A4 mAb using Bio-layer interferometry (BLI) Octet instrument

For antibody affinity measurement, antibody (30 µg/ml) was loaded onto the protein A biosensors (Sartorius, USA) for 4 min. Following a short baseline kinetics buffer, the loaded biosensors were incubated with recombinant S100A4 protein (HIS tagged, Sino Biological) at a series of concentration titrations. Kinetic sensor grams for each antibody are collected and background was corrected using a reference well without analyte for sensor drifting, ForteBio's data analysis software was used to fit the data to a 1:1 binding model and $K_D$ was calculated using the ratio of dissociation rate ($k_{off}$)/ and an association rate ($k_{on}$).

##### Measurement of anti-S100A4 mAb binding using ELISA

Binding to S100A4 protein by antibodies from B cell culture supernatants or purified mAbs was determined by ELISA. Antibody concentration titration was used to estimate binding affinity ($E_{50}$) using a 4-parameter fitting model with GraphPad Prism software (version 8).

##### Chick DRG neurite growth assay

Chick DRGs from E7 embryonated egg was isolated in cold PBS and individually placed on Nucleopore membranes (SIGMA Millipore). Premixed collagen/DMEM medium supplemented with recombinant mouse S100A4 protein (25 µg/ml, Sinobiological) with or without S100A4 antibodies (200 μg/ml) were placed over the DRGs, triplicates per condition. After incubation for 3 days, DRGs were fixed and stained with anti- ß-III-tubulin antibody and Alex-594 secondary antibody. Each DRG was imaged by Zeiss Axio Imager 2 with apotome 2.0, and neurite outgrowth was measured and compared in Image J software.

**Pharmacokinetic evaluation of S100A4 antibody**

Groups of C57BL/6 mice (5 mice/group) were administered with 20 mg/kg of antibody via intraperitoneal injection. Blood and tissue samples were at 4 h, 1, 2, 4, and 7 days. Serum samples were obtained from blood after incubation at room temperature for 20 minutes and centrifugation at 4000g for 10 minutes. Tissue samples were lysed by NP40 solution (GenDepot). For the antigen-binding ELISA, 96-well high binding plates (Corning Inc, Corning, NY) were coated with 0.2µg/100µL S100A4 antibody in PBS at 4°C overnight. After coating, the plates were washed with PBST (0.05% Tween20) and blocked with 3%BSA in PBS for 1h at room temperature. Samples were loaded into the wells after appropriate dilutions with Assay Diluent Buffer (Biolegend, San Diego, CA) and incubated for 1 h at room temperature under shaking. After washing, goat anti rabbit IgG-HRP (Cell Signaling, Danvers, MA) was added and incubated for 30 minutes at room temperature. After being washed three times by TBST, each well was incubated with 100µL of 1-Step Turbo TMB ELISA substrate (Thermo Fisher, Rockford, IL) for 10 minutes at room temperature. The reaction was stopped with TMB stop solution (Biolegend, San Diego, CA) and the plates were read at 450nm.

**References**


76. X. Gui *et al.*, Disrupting LILRB4/APOE Interaction by an Efficacious Humanized Antibody Reverses T-cell Suppression and Blocks AML Development. *Cancer Immunol Res* **7**, 1244-1257 (2019).

77. P. Zhao *et al.*, Enhanced anti-angiogenetic effect of transferrin receptor-mediated delivery of VEGF-trap in a glioblastoma mouse model. *MAbs* **14**, 2057269 (2022).
78. J. Liu *et al.*, A highly selective humanized DDR1 mAb reverses immune exclusion by disrupting collagen fiber alignment in breast cancer. *J Immunother Cancer* **11**, (2023).

**Supplementary Figure 1**

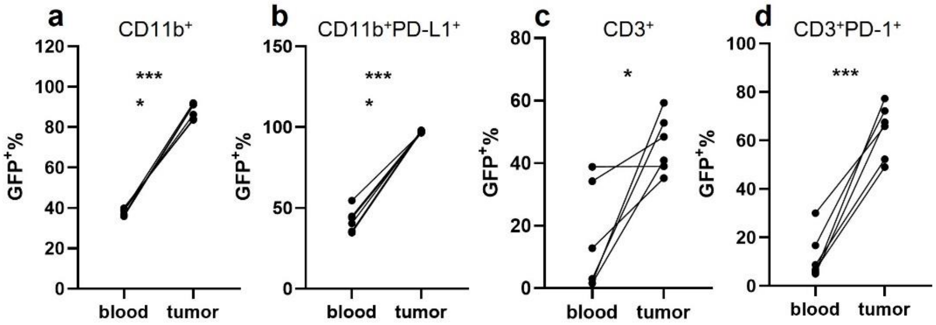


**Supplementary Fig 1. S100A4$^+$/GFP$^+$ expression comparison between blood and tumor stromal cells in the E0771 tumor-bearing *S100a4$^{+/-}$* mice**

Compared with blood samples, S100A$^+$/GFP$^+$ expression level is significantly higher in the CD11b$^+$ (a), CD11b$^+$PDL1$^+$ (b), CD3$^+$ (c), and CD3$^+$PD-1$^+$ (d) cells in the E0771 tumor microenvironment of *S100a4$^{+/-}$* mice. Paired *t*-test analysis was used. * $p$<0.05; *** $p$<0.001; **** $p$<0.0001.

### Supplementary Figure 2

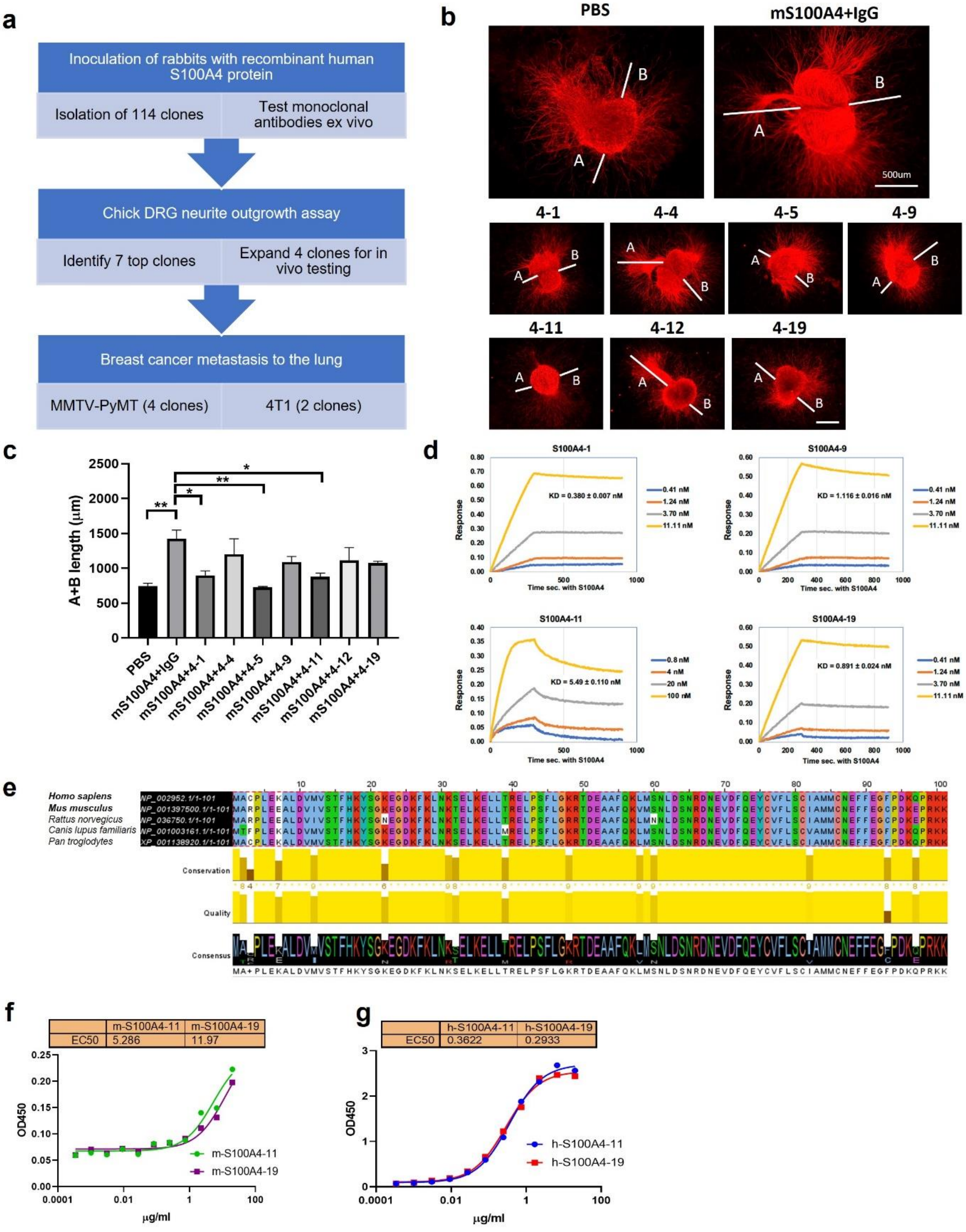

**Supplementary Fig 2. Anti-human S100A4 antibody screen design and characterization**

(a) A schematic summary of S100A4 antibody screen. (b) Dorsal root ganglia (DRG) were harvested from 7-day-old chick embryos. They were cultured for 3 days with soluble S100A4 protein plus IgG or one of 114 monoclonal S100A4 antibodies against human S100A4 protein in collagen matrix. DRG were fixed and stained with β3-Tubulin antibody to measure the lengths of the neurite outgrowth (A+B). Scale bar denotes 500 µm. (c) S100A4-1, 4-5, and 4-11 clones significantly suppressed S100A4-induced neurite outgrowth. One-way ANOVA analysis was used. *$p<0.05$; **$p<0.01$. (d) Kinetic binding sensorgrams of S100A4 mAbs are shown and concentrations of S100A4 (as analyte) used for fitting and calculation of $K_D$ are indicated in each graph. Each mAb name is indicated on the top of each graph. (e) Sequence alignments of S100A4 proteins (by Jalview 2.11.2.6) from *Homo sapiens* (human), *Mus musculus* (mouse), *Rattus norvegicus* (rat), *Canis lupus familiaris* (dog), and *Pan troglodytes* (chimpanzee), whose sequence data were collected from NCBI. The S100A4 protein is highly conserved among species. (f, g) S100A4-11 and S100A4-19 antibody binding affinities to mouse (f) and human (g) S100A4 proteins. ELISA concentration titration assay results show that 4-11 antibody has 2x fold affinity to mouse S100A4 protein, compared with 4-19. S100A4-11 antibody has ~10-fold higher affinity for the human S100A4 protein than mouse protein.

**Supplementary Figure 3**

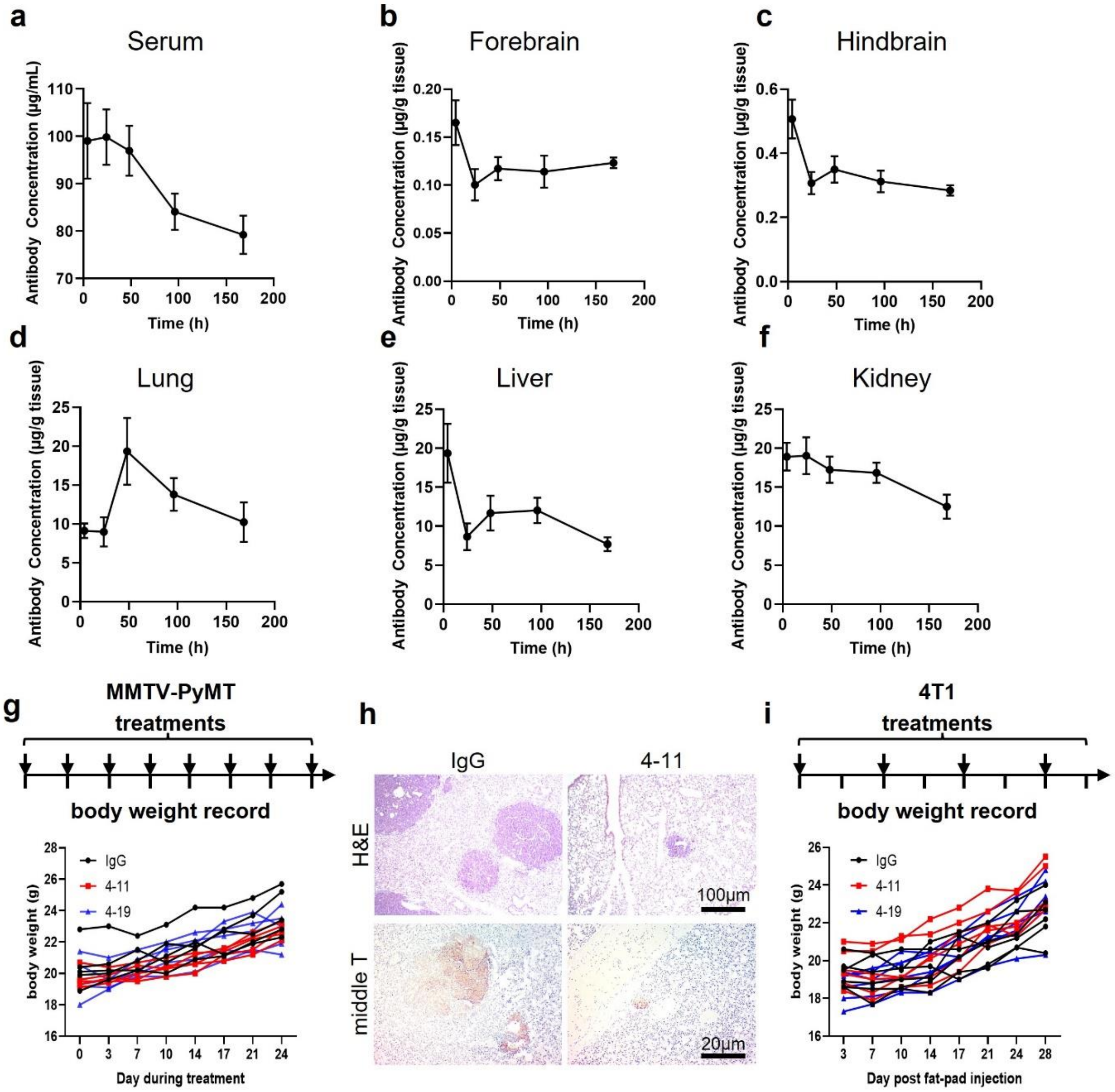


**Supplementary Figure 3. Pharmacokinetic profile and tolerance of S100A4-11 antibody**

(a) serum (b) forebrain, (c) hindbrain, (d) lung, (e) liver, and (f) kidney from B6 mice were harvested at 4, 24, 48, 96, and 168 hr post S100A4-11 treatment (20mg/kg, IP). Each time point has 5 mice. Error bar is mean ± SEM. (g) Body weight measurement of IgG, 4-11 or 4-19 treated MMTV-PyMT mice (n=5 per group). (h) Hematoxylin and eosin staining and polyomavirus middle T antibody staining in lung metastatic nodules from representative IgG control and 4-11 antibody treated mice. (i) Body weight measurement of IgG, 4-11 or 4-19 treated 4T1 tumor bearing mice (n=6 per group). Body weights were recorded twice per week.

**Supplementary Figure 4**

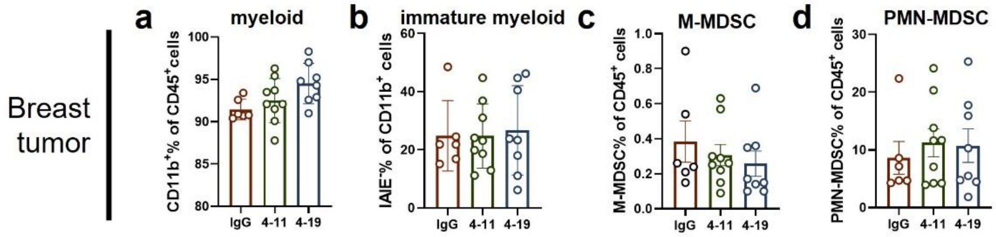


**Supplementary Figure 4. Effects of S100A4-11 antibody treatment on the MDSCs in the MMTV-PyMT primary breast tumor**

Flow cytometry analysis of primary breast tumors showed no significant difference in the total myeloid (a), immature myeloid (b), PMN-MDSC (c), and M-MDSC (d). n=6-9 per group.

**Supplementary Figure 5**

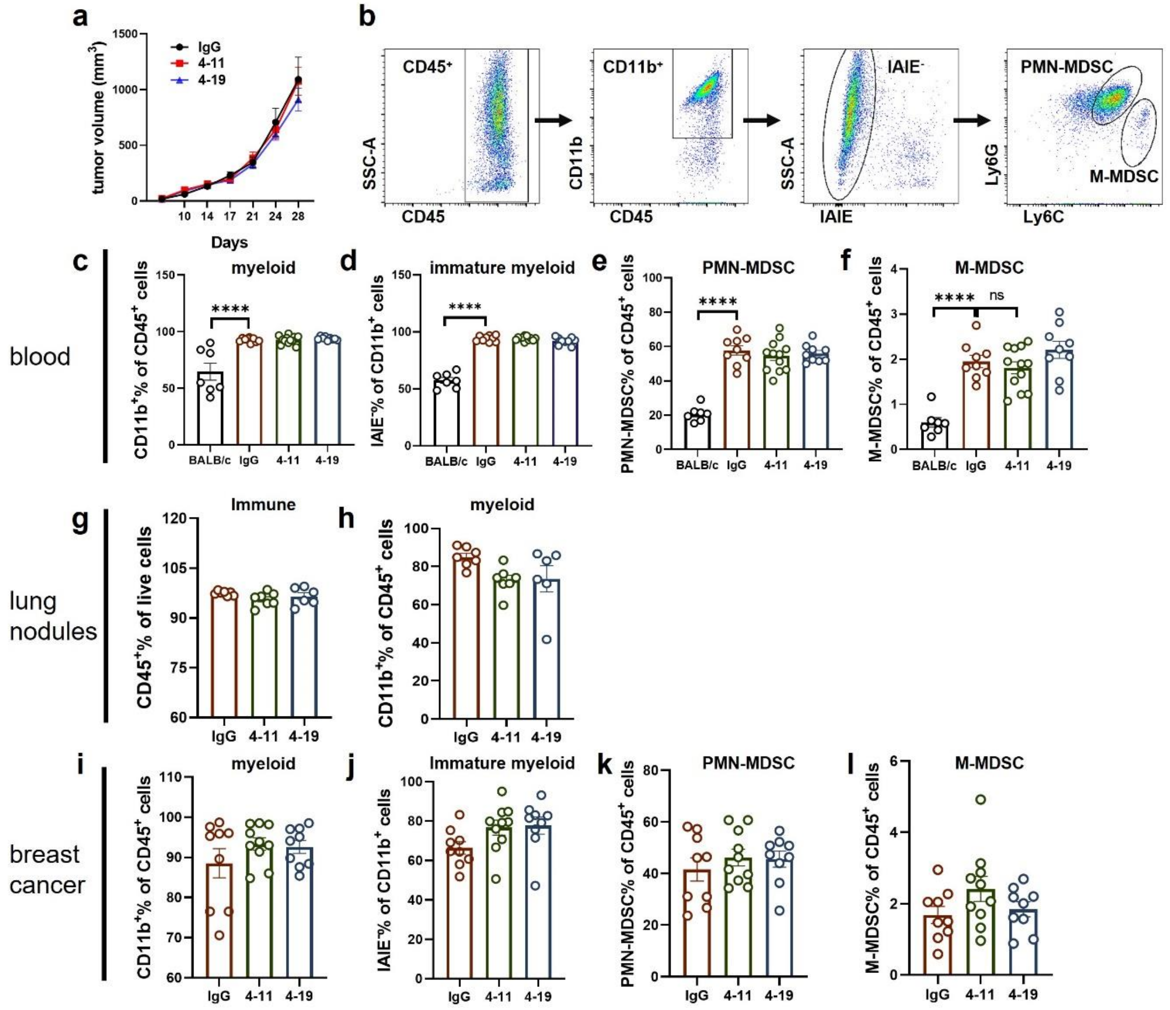


**Supplementary Figure 5. Effects of S100A4-11 antibody treatment on the blood and tumor MDSCs in 4T1 breast cancer model**

(a) The 4T1 breast tumor volumes were measured twice a week for IgG, 4-11 and 4-19 treated mice (n=5-6). (b) Flow cytometry gating strategy of blood samples after discriminating doublets and dead cells. (c-l) Flow cytometry analysis of PBMCs: (c-f) myeloid cells (c) and immature myeloid cells (d), PMN-MDSCs (e) and M-MDSCs (f) in the blood. Tumor bearing mice (n=9-12 per treatment group), non-tumor bearing control BALB/c mice (n=7). (g, h) The total immune cells (g), and myeloid cells (h) in lungs of treated mice. (n=6-7) (i-l) Fractions of myeloid cells (i),

IAIE/MHC2-negative myeloid cells (j), PMN-MDSC (k), and M-MDSC (l) in the breast tumor (n=9-10 per treatment group). Ordinary one-way ANOVA analysis was used. Error bar indicates mean ± SEM. ****$p$<0.0001; ns, not significant.

**Supplementary Figure 6**

a

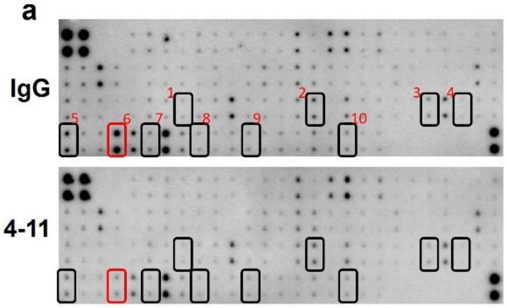


b

| # | Protein |
|---|---|
| 1 | IL-20 |
| 2 | Lungkine/CXCL15 |
| 3 | MIP-1 gamma (CCL9) |
| 4 | MIP-3 alpha (CCL20) |
| 5 | Osteopontin (OPN) |
| 6 | Pro-MMP9 |
| 7 | SCF |
| 8 | TACI |
| 9 | TPO |
| 10 | VEGF R1 |

**Supplementary Fig 6. Cytokine array analysis of lung lysates from 4T1 tumor-bearing mice**

(a) Lung tissue lysates were harvested from 4T1 tumor-bearing mice on day 14 post-injection and applied to Raybiotech C6 mouse cytokine array. Representative images were taken by chemiluminescence exposure. After quantitative analysis, 10 proteins (square) were identified to be downregulated in the lung tissue of 4-11 treated mice. Pro-MMP9 is labeled by a red box. Cytokine array image shown is representative from two independent experiments. (b) 10 proteins that showed consistent and significant changes in IgG vs 4-11 treated mice in two different experiments.

#### Supplementary Figure 7

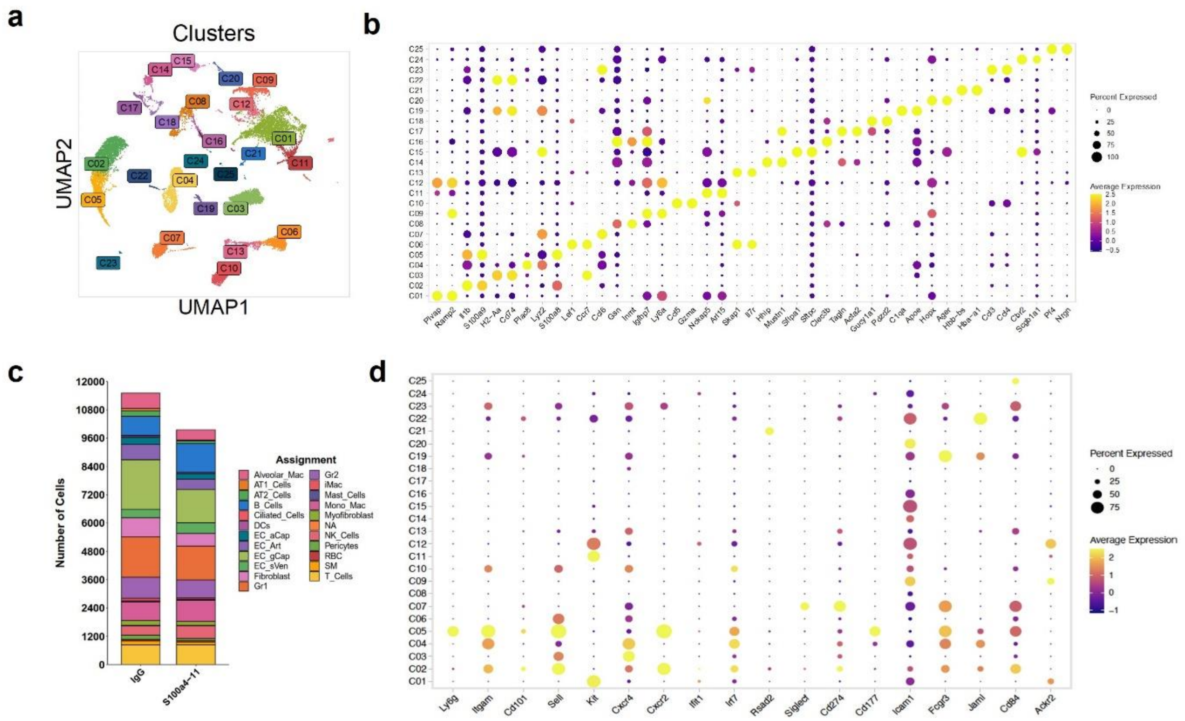


**Supplementary Figure 7. Cell type assignments of single-cell RNA-sequencing clusters**

(a) Cell clustering was performed on lung tissue samples from IgG and 4-11 treated mice. UMAP projection of all cells, color coded by assignment shown in (c). (b) Dot plot showing top2 differentially expressed gene by cluster. (c) Bar graphs showing numbers of different cell types present in the lungs from IgG and S100A4-11 antibody-treated mice. (d) Dot plot showing neutrophil subtype markers.

### Supplementary Figure 8

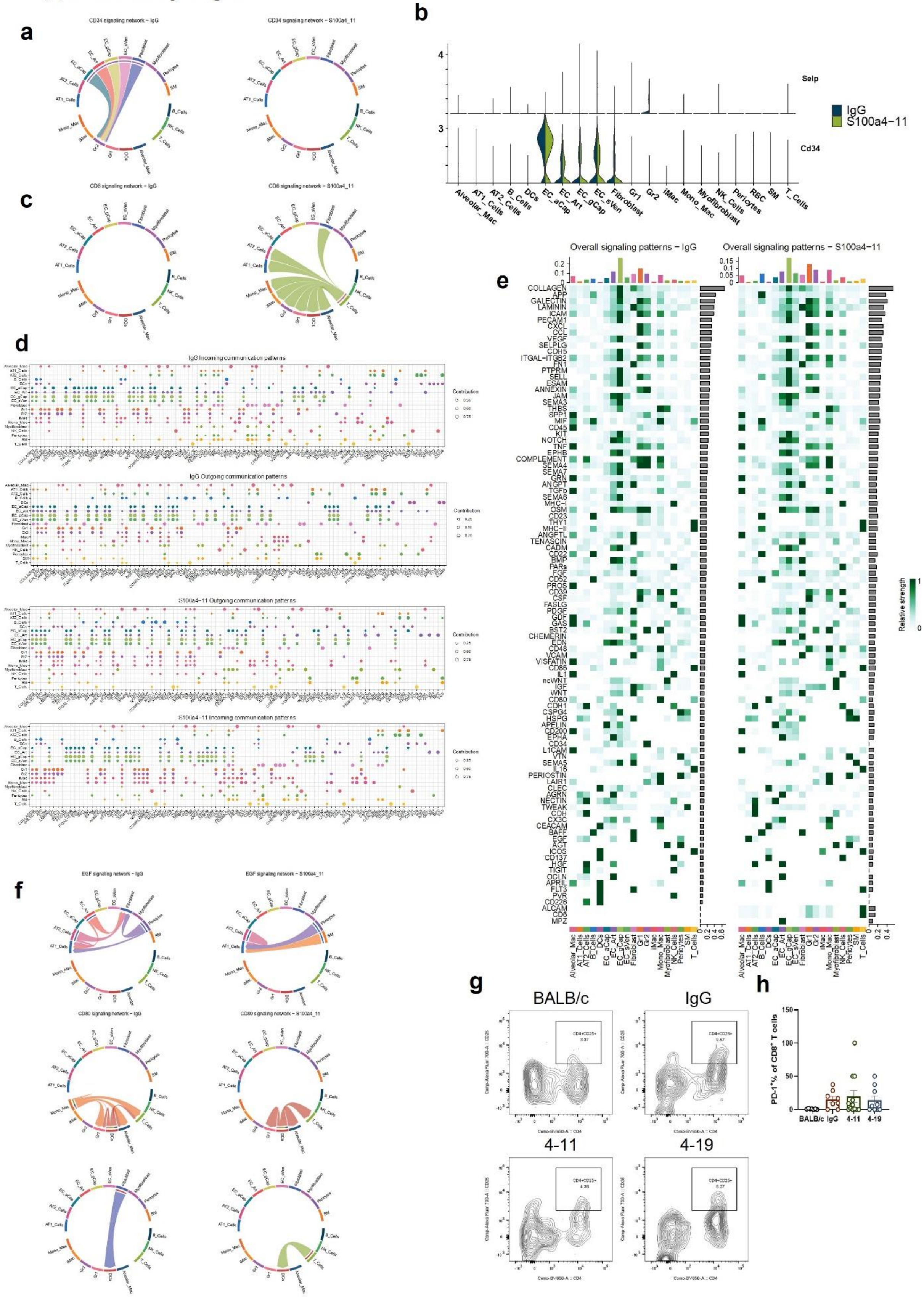

**Supplementary Figure 8. Differential cell:cell communications among various cell types in the pre-metastatic lung** (a) Chord diagram showing CD34 signaling network strengths between lung from IgG vs 4-11 treated mice. (b) Violin plot showing expression of *Selp* and *CD34* in lung cells (day 14). (d) Dot plots showing overall incoming/outgoing enriched signaling pathways from different cell types and molecules between IgG vs 4-11 treated group. (e) The Overall signaling patterns among lung cells in IgG vs. S1004-11 treated group. (f) Chord diagram showing EGF, FLT3, and CD80 signaling network strengths between IgG vs 4-11 treated group. (g) Flow cytometry analysis of blood for $CD4^{+}CD25^{+}$ Treg cells. (h) Flow cytometry analysis of blood showing the percentage of $PD\text{-}1^{+}/CD8^{+}$ T cells (BALB/c: n=7; tumor-bearing group: n=8-12). Ordinary one-way ANOVA analysis was used. Error bar denotes mean ± SEM. *$p<0.05$.

#### Supplementary Figure 9

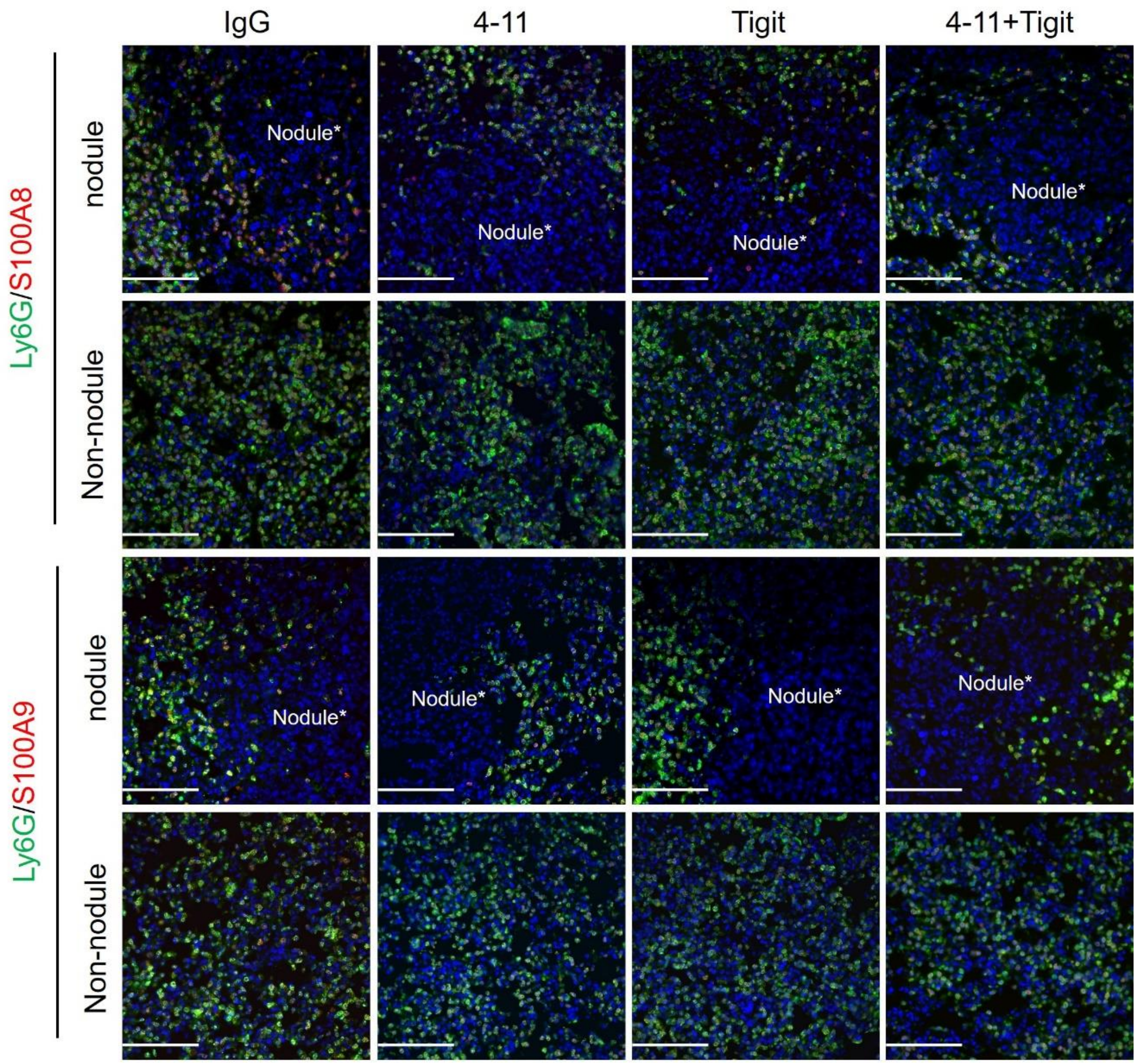


**Supplementary Figure 9. S100A4-11 plus Tigit antibody treatment has no effect on the number of infiltrating neutrophils in late-stage metastatic lung**

Late-stage metastatic lung tissue stained for Ly6G/S100A8 or S100A9 (n=3). Scale bar denotes 100µm.

**Supplementary Table1: The antibodies list used for IHC/IF and western blotting**

| Antibody | Cat# | Concentration | Company |
|---|---|---|---|
| CD163 | MA5-11458 | 1:200 | ThermoFisher |
| CD206 | MCA2235 | 1:200 | Bio-Rad |
| Pan-Keratin | 26411-1-AP | 1:200 | Proteintech |
| S100A4 | #13018 | 1:200 | Cell Signaling |
| S100A8 | #47310 | 1:200 | Cell Signaling |
| S100A9 | #73425 | 1:200 | Cell Signaling |
| Ly6g | 127602 | 1:200 | Biolegend |
| Polyome virus, Medium T | NB100-2749 | 1:50 | Novus Biology |
| MMP9 | ab76003 | 1:200 | Abcam |
| NKp46 | AF2225 | 1:250 | R&D system |
| CD8α | 14-0915-82 | 1:250 | eBioscience |
| Perforin | #31647 | 1:250 | Cell Signaling |
| Tigit | ab300073 | 1:250 | abcam |
| MMP9 | PA5-13199 | 1:1,1000 (WB) | ThermoFisher |
| beta Actin | A00730 | 1:3,000-5,000 (WB) | Genscript |

**Supplementary Table2: The antibodies list used for flow cytometry**

| Antibody | Cat# | Fluorescence | Company |
|---|---|---|---|
| CD45 | 103114 | PE-Cy7 | Biolegend |
| CD45 | 103116 | APC-Cy7 | Biolegend |
| CD11b | 101208 | PE | Biolegend |
| CD11b | 101216 | PE-Cy7 | Biolegend |
| CD11b | 101259 | BV650 | Biolegend |
| I-A/I-E (MHC-II) | 107614 | APC | Biolegend |
| I-A/I-E (MHC-II) | 107628 | APC-Cy7 | Biolegend |
| Ly6c | 128037 | BV711 | Biolegend |
| Ly6g | 127624 | APC-Cy7 | Biolegend |
| CD206 | 141723 | BV650 | Biolegend |
| CD163 | 156704 | PE | Biolegend |
| CD3 | 100308 | PE | Biolegend |
| CD4 | 100469 | BV650 | Biolegend |
| CD8a | 100747 | BV711 | Biolegend |
| Tigit | 156105 | APC | Biolegend |
| PD-1 | 135216 | PE-Cy7 | Biolegend |
| CD69 | 104539 | AF700 | Biolegend |
| CD25 | 102024 | AF700 | Biolegend |
